# Comparative multi-omics analyses of cardiac mitochondrial stress in three mouse models of frataxin deficiency

**DOI:** 10.1101/2023.02.03.526305

**Authors:** Nicole M. Sayles, Jill S. Napierala, Josef Anrather, Nadège Diedhiou, Jixue Li, Marek Napierala, Hélène Puccio, Giovanni Manfredi

## Abstract

Cardiomyopathy is often fatal in Friedreich Ataxia (FA). However, the FA heart maintains adequate function until disease end stage, suggesting that it can initially adapt to the loss of frataxin (FXN). Conditional knockout mouse models with no *Fxn* expression show transcriptional and metabolic profiles of cardiomyopathy and mitochondrial integrated stress response (ISR^mt^). However, ISR^mt^ has not been investigated in models with disease-relevant, partial decrease of FXN. We characterized the heart transcriptomes and metabolomes of three mouse models of partial FXN loss, YG8-800, KIKO-700, and Fxn^G127V^. Few metabolites were significantly changed in YG8-800 mice and did not provide a signature of cardiomyopathy or ISR^mt^. Instead, several metabolites were altered in Fxn^G127V^ and KIKO-700 hearts. Transcriptional changes were found in all models, but differentially expressed genes consistent with cardiomyopathy and ISR^mt^ were only identified in Fxn^G127V^ hearts. However, these changes were surprisingly mild even at an advanced age (18-months), despite a severe decrease in FXN levels to 1% of WT. These findings indicate that the mouse heart has extremely low reliance on FXN, highlighting the difficulty in modeling genetically relevant FA cardiomyopathy.

**Summary statement:** The mitochondrial integrated stress response in the heart of a Friedreich Ataxia mouse model is surprisingly mild, despite a severe decrease in frataxin levels below 1% of normal.

## Introduction

Cardiomyopathy is one of the most prominent pathological features of Friedreich Ataxia (FA), a genetic mitochondrial disease caused by decreased levels of frataxin (FXN) protein resulting in impaired iron-sulfur (Fe-S) cluster biogenesis. Heart failure is the cause of death in ~60% of patients with FA (Payne, 2022), which clearly makes cardiomyopathy a primary target for therapy. FXN deficiency is most commonly due to a trinucleotide repeat expansion (GAA) in the first intron of the *Fxn* gene (Campuzano et al., 1996). Rarely, FA is associated with compound heterozygosity with one expanded allele and one allele harboring an exonic mutation. A G130V substitution has been associated with forms of FA with milder clinical presentation and slower progression (Galea et al., 2016). Pathologically, FA patients develop a hypertrophic cardiomyopathy, characterized by a profound disarray of cardiomyocytes cytological architecture, with mitochondrial proliferation and iron deposition (Hanson et al., 2019).

FA patients with cardiomyopathy display impaired myocardial perfusion reserve index associated with microvascular alterations and tissue fibrosis. Arrhythmias are also common and can contribute to mortality (Bidichandani and Delatycki, 1993; Koeppen, 2011). Typically, death from cardiomyopathy in FA occurs in the third or fourth decade of life. Surprisingly, the FA heart often maintains adequate systolic function until shortly before death, even though the underlying causes of tissue degeneration, oxidative phosphorylation (OXPHOS) dysfunction, impaired iron homeostasis, and oxidative stress are likely present early on, possibly even during development (Palau, 2001). This suggests that the FA heart can adapt, at least initially, to the OXPHOS defects caused by the loss of FXN. This adaptation likely involves metabolic rewiring to allow the utilization of alternative energy sources, which do not depend on OXPHOS.

The normal heart relies heavily on fatty acids as the main energy source (Lopaschuk et al., 2010). However, the enzymatic steps required to generate ATP from fatty acids are defective in FA, due to Fe-S cluster deficiency (Rotig et al., 1997). Therefore, the FA heart shifts its metabolism towards aerobic glycolysis (Payne and Wagner, 2012). This process is bioenergetically less efficient compared to mitochondrial OXPHOS, but it could be tolerated, if enough glucose is available for glycolysis. In parallel to this metabolic adjustment, several other adaptive events have been demonstrated to take place in the heart of a mouse model of cardiac *Fxn* deletion. These include a paradoxical iron-starvation response driven by the iron response protein 1, which senses the depletion of Fe-S clusters (Martelli and Puccio, 2014). This response contributes to the accumulation of mitochondrial iron (Martelli et al., 2015), which is also a key pathological feature of FA cardiomyopathy in human patients (Lamarche et al., 1980). Furthermore, there is accumulation of unutilized lipids in cardiomyocytes. There is also a build-up of oxidative stress, presumably initiated in mitochondria. The consequences of oxidative stress may not be immediately apparent, because they are matched by a strong upregulation of antioxidant defenses. This complex metabolic adaptation occurring in the FA heart recapitulates the main metabolic features of a process that has been well characterized in mitochondrial diseases, defined as mitochondrial integrated stress response (ISR^mt^) (Dogan et al., 2014; Forsstrom et al., 2019; Kaspar et al., 2021; Khan et al., 2017; Kuhl et al., 2017; Nikkanen et al., 2016; Sayles et al., 2022).

The ISR^mt^ can occur as the result of protein misfolding in mitochondria (Narayana Rao et al., 2022), like the endoplasmic reticulum (ER) unfolded protein response leading to ER stress. Both the ISR^mt^ and ER stress converge onto the activation of the transcription factor ATF4, which downregulates the transcription of many genes, while upregulating expression of genes encoding chaperones, proteases, proteasome subunits, and other factors involved in proteostasis. However, the ISR^mt^ has also an important mitochondrial arm, driven by another transcription factor, ATF5, which downregulates the expression of genes encoding mitochondrial proteins, including respiratory chain subunits, transporters, carriers, mitochondrial DNA replication factors, while activating the expression of genes encoding mitochondrial chaperones and proteases, aimed to reduce mitochondrial proteotoxic stress.

The ISR^mt^ also involves a profound rewiring of cellular metabolism, including activation of Akt and mTOR signaling (Aramburu et al., 2014; Wu and Storey, 2021) and upregulation of serine one-carbon (1C) metabolism, a well-established metabolic consequence of the ISR^mt^. The serine-1C metabolic pathway provides one carbon units for methionine biosynthesis, using folate carriers (Mehrmohamadi et al., 2014; Nikkanen et al., 2016), and facilitates glutathione (GSH) production to antagonize oxidative stress. At the same time, it allows for enhanced production of NADPH, which is necessary to reduce oxidized GSH (GSSG). However, while serine can be derived from the diet, under ISR^mt^, increased serine-1C metabolism can force the utilization of glucose for de novo serine biosynthesis, thereby decreasing the glucose available for energy generation by glycolysis and other metabolic pathways, such as nucleotide biosynthesis.

The ISR^mt^ has also significant inter-organ metabolic effects. Notably, it stimulates the production in heart and skeletal muscle of myokines, such as GDF15 and FGF21, which are secreted in the blood stream and signal to the liver and the adipose tissue to activate gluconeogenesis and fatty acid mobilization, respectively (Boenzi and Diodato, 2018). This can lead to loss of fat stores and accumulation of lipids in heart and muscle, which cannot oxidize them due to the downregulation of mitochondrial □-oxidation associated with ISR^mt^ (Forsstrom et al., 2019).

There is evidence that ISR^mt^ activation occurs in the hearts of mouse models of *Fxn* genetic ablation. In a heart-specific mouse model with conditional *Fxn* knockout (cKO), there was elevation of ATF4 accompanied by suppression of protein synthesis, elevation of chaperones and proteases, and upregulation of serine 1C-metabolism (Huang et al., 2013). These mice also exhibit increased phosphorylation of AKT and dysregulation of multiple downstream effectors of mTORC1 (Tong et al., 2022). Furthermore, serine-1C metabolism elevation and FGF21 upregulation were described in the heart of a mouse model with inducible silencing of *Fxn* (Vasquez-Trincado et al., 2021). However, these studies examined mouse models that do not fully reflect the physiology of FA in humans, because the complete genetic ablation of *Fxn* in the heart or the acute postnatal silencing of *Fxn* differ from the situation in the human heart, where the defect is present since embryonic development and FXN is never completely absent. Investigating ISR^mt^ and a clearer understanding of its metabolic consequences in a “humanized” mouse model of FA could help to determine if modulation of this pathway would be beneficial for treating FA cardiomyopathy. Therefore, to assess the effects of heart FXN deficiency in more physiological model systems, we investigated three independent FA mouse models of partial FXN loss using an integrated multi-omics approach. We performed unbiased transcriptomics and metabolomics analyses of hearts from aged (18-months) mice that model partial FXN deficiency, due to either intron 1 GAA expansion in the endogenous *Fxn* gene or to the expression of a human GAA expanded *Fxn* transgene in a mouse *Fxn* KO background or to a homozygote knock in (KI) *Fxn* missense mutation (G127V), the mouse equivalent of the human pathogenic mutation (G130V) (Bidichandani et al., 1997; Galea et al., 2016). Notably, none of these FXN deficient models have premature death, at least before age 18 months, suggesting that if cardiomyopathy is present, it is mild, unlike the cKO or acute silencing models that cause rapidly fatal cardiomyopathy.

## Results

### Gene expression profiles indicate mitochondrial and cardiac stress in the YG8-800 mouse

To investigate if heart ISR^mt^ and related metabolic rewiring occur in a mouse model of partial FXN deficiency, we performed unbiased metabolomics and transcriptomics analyses in the heart of a humanized FA model, the Tg(FXN)YG8Pook/800J (YG8-800) mouse from The Jackson Laboratories. YG8-800 transgenic mice are homozygous for a *Fxn* constitutive null allele (exon 2 deletion) and hemizygous for a human *Fxn* transgene, which contains >800 GAA repeats in intron 1. Recently, the heart of this mouse model was characterized, and by ELISA the levels of cardiac human FXN were estimated to be approximately 5% relative to a control mouse expressing human *FXN* with only 9 GAA repeats (Gerard et al., 2023). Cardiac involvement included increased heart to body weight ratio and moderately reduced ejection fraction at 6 months of age. At this time, the cardiac phenotype was defined as being at an early stage of progression. Thus, we opted to investigate the metabolomic and transcriptomic profiles of YG8-800 hearts and littermate controls (WT), expressing normal levels of mouse FXN and no human FXN, at 18 months of age, expecting that cardiac alterations would be more severe in the aging mouse. By ELISA assay, we find that the YG8-800 mice expressed lower levels of FXN (21±4 ng/mg) compared to endogenous mouse levels in the WT littermate controls (208 ±16 ng/mg) (Fig. S1A). Since no differences were reported between males and females (Gerard et al., 2023), we utilized only males for these studies.

Hierarchical cluster analysis of metabolomics data did not show clustering of samples by genotype (Fig. 1A), indicating there were very few metabolic differences between YG8-800 and WT hearts, with only 6 metabolites (4-pyridoxic acid, NADH, kynurenic acid, glutathione, deoxyribose 5-phosphate, and guanosine triphosphate) reaching a significance threshold (p < 0.05). Unbiased metabolic Kyoto Encyclopedia of Genes and Genomes (KEGG) pathway analysis by MetaboAnalyst on these few metabolites suggested alterations of vitamin B6 metabolism (Fig. 1B). On the other hand, transcriptomic analysis of YG8-800 hearts showed several differentially expressed genes (DEGs; with a threshold set at p < 0.05) (1052 DEGs), the majority of which were downregulated (669 DEGs downregulated and 383 upregulated) (Fig. 1C). Unbiased gene ontology (GO) analyses on all DEGs (p < 0.05) revealed several significantly altered pathways. The top 10 most significant upregulated pathways were broad categories that did not suggest cardiac stress (Fig. 1D). However, among the top 10 most significant downregulated pathways, there were several enrichments of interest, including protein translation, ribosome biogenesis, mitochondrion organization, and protein import into mitochondrial matrix (Fig. 1E).

**Figure 1.**
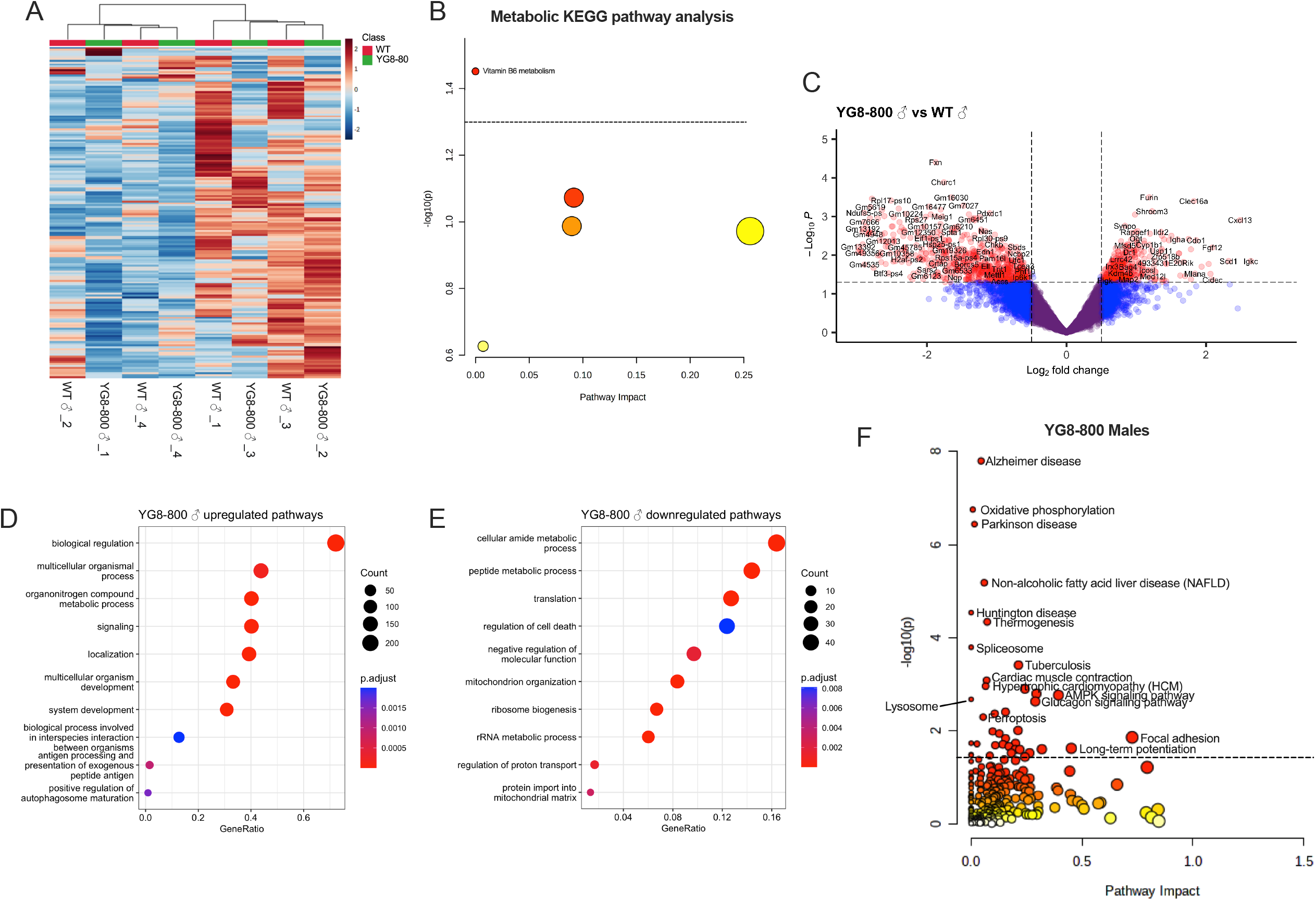
Enrichment of pathways associated with mitochondrial and cardiac stress in YG8-800. (A) Heatmap of all measured metabolites in WT and YG8-800 hearts (n = 4 18-months old mice). Red indicates increased abundance and blue indicates reduced abundance. (B) Unbiased metabolic KEGG pathway analysis. Pathway impact is a combination of the centrality and pathway enrichment results, calculated as the sum of the importance measures of each of the matched metabolites and then divided by the sum of the importance measures of all metabolites in each pathway. For each data point, the size of the circle is correlated with pathway impact and the deeper color indicates higher significance. (C) Volcano plot of gene expression (n = 4 18-months old mice). Genes in red indicate p > 0.05, FC > 0.5. (E, F) Top 10 results of unbiased gene ontology analyses of significantly upregulated (D) or downregulated (E) genes. Gene Ratio is calculated by dividing the number of genes that belong to a given gene set by the total number of genes in the gene set. Count indicated by size and adjusted p value (p.adjust) indicated by color. (F) Integrated metabolomics and transcriptomics data from YG8-800 males. KEGG IDs of some significantly enriched pathways (−Log_10_(p) > 1.3) were added to the joint pathway analysis in MetaboAnalyst.

Next, we looked at the integrated pathways with MetaboAnalyst joint-pathway analysis of the transcriptomic and metabolomic data. An initial analysis including ribosomal genes resulted in an overrepresentation of the ribosome and RNA transport pathways (Fig. S1B), which hindered the visualization of other enriched pathways. Therefore, we performed another integrated analysis excluding ribosomal subunit genes that modified the significance of the remaining pathways and allowed for a better visualization of several pathways related to hypertrophic cardiomyopathy, ferroptosis, and oxidative phosphorylation, in addition to pathways associated with neurodegeneration (Fig. 1F). Overall, these results suggest cardiac stress without evidence of ISR^mt^.

### KIKO-700 mice display metabolic and transcriptional alterations associated with cardiac stress and ISR^mt^

Next, we analyzed another model of partial FXN deficiency, a knock-in, knock-out (KIKO) mouse model. In this model, one allele of mouse *Fxn is* deleted, while the other allele harbors a GAA expansion in intron 1. In a well-established FA KIKO mouse model harboring >230 GAA repeats (KIKO-230), approximately 30% residual FXN protein is detected in the heart (Miranda et al., 2002). This level of residual FXN was shown to be sufficient to prevent iron accumulation and severe fibrosis in the heart (Miranda et al., 2002). Herein, we investigated the cardiac effects of a newly generated KIKO mouse with a longer GAA expansion, harboring 700 repeats (KIKO-700). Longer expansions in FA patients are associated with earlier disease onset, increased disease severity, and left ventricular hypertrophy (Isnard et al., 1997; Koeppen, 2011). Surprisingly, KIKO-700 hearts at 18 months of age had only a 50% reduction in FXN protein as estimated by western blot normalized by total protein (Fig. 2A and S2A). Therefore, despite the longer GAA expansion in the KIKO-700 mice, the levels of residual heart FXN are higher than those reported for the KIKO-230 model.

**Figure 2.**
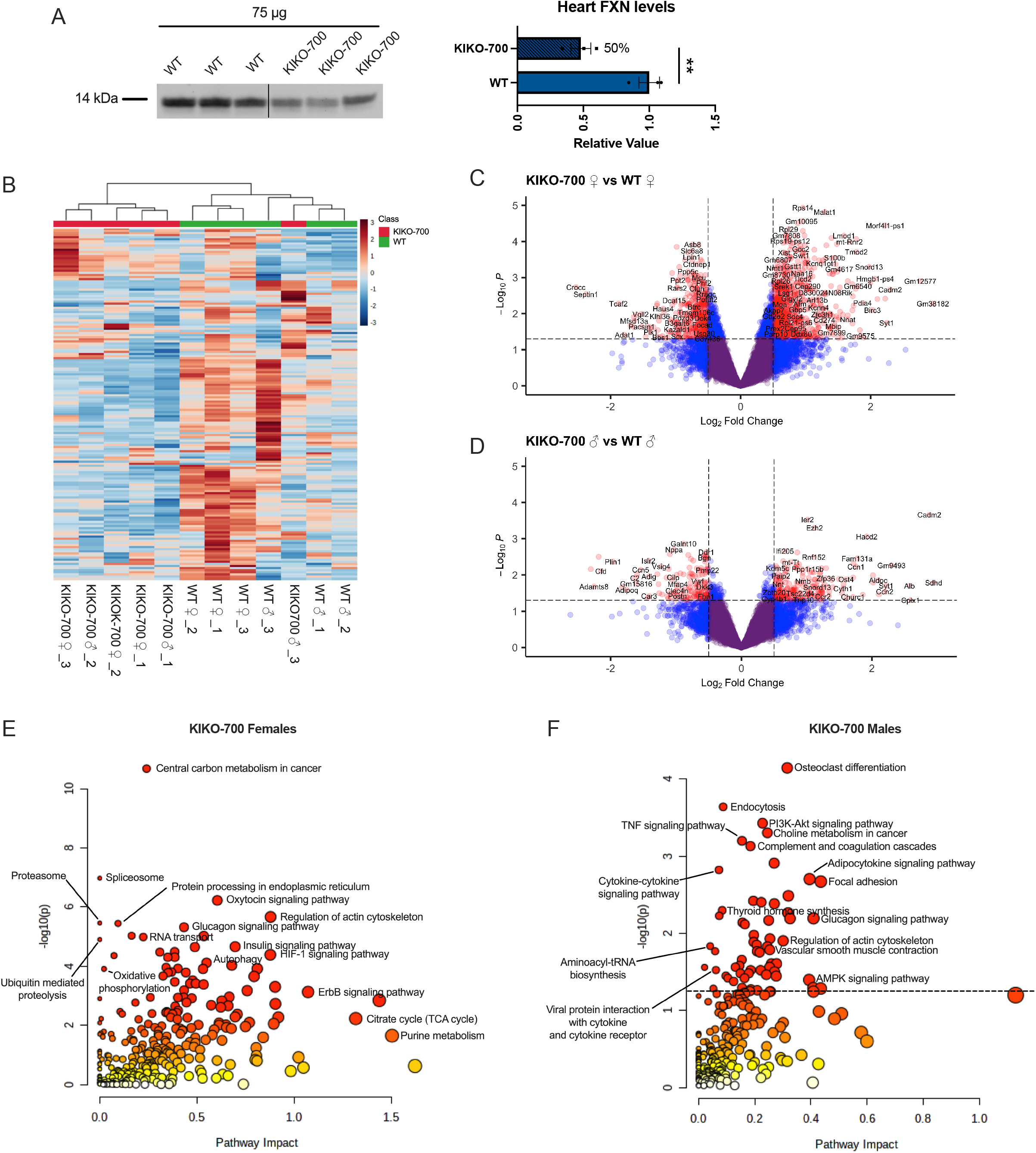
Enrichment of pathways associated with ISR^mt^ and cardiac stress in KIKO-700. (A) Western blot analysis of FXN expression in WT and KIKO-700 male hearts, quantified in the right panel (n = 3) (statistical analysis was performed with unpaired two-tailed *t*-test, ** = p < 0.009, data represented as mean ± SEM). (B) Heatmap of all measured metabolites in WT and KIKO-700 hearts (n = 3/sex/genotype 18-months old mice). Red indicates increased abundance and blue indicates reduced abundance. (C, D) Volcano plot of gene expression in KIKO-700 females (C) and males (D) (n = 3/sex/genotype 18-months old mice). Genes in red indicate p > 0.05, FC > 0.5. (E, F) Integrated metabolomics and transcriptomics data from KIKO-700 females (E) and males (F).

Metabolomics and transcriptomics analyses were performed on both male and female KIKO-700 and WT hearts. Hierarchical clustering of detected metabolites revealed partial clustering by genotype (Fig. 2B). To note, most differential metabolites in KIKO-700 hearts (p < 0.05) were decreased in abundance. Unbiased KEGG metabolic pathway analysis revealed several enriched pathways. In female KIKO-700 hearts, glutathione metabolism, purine metabolism, citric acid cycle (Tricarboxylic acid cycle), and several amino acid-related metabolic pathways were significantly enriched (Fig. S3A). Although male KIKO-700 hearts had fewer differential metabolites than female hearts (15 vs. 58 metabolites), some of the same pathways were enriched, including purine metabolism, arginine biosynthesis, and pentose phosphate (Fig. S3B). Of these pathways, glutathione metabolism and nucleotide biosynthesis have been associated with ISR^mt^ and found to be dysregulated in various mouse models of mitochondrial dysfunction and hypertrophic cardiomyopathy (Dogan et al., 2014; Forsstrom et al., 2019; Kaspar et al., 2021; Khan et al., 2017; Kuhl et al., 2017; Nikkanen et al., 2016; Sayles et al., 2022). Furthermore, several DEGs were identified in KIKO-700 hearts, with a higher number of DEGs in females (1804 DEGs in females and 505 DEGs in males) (Fig. 2C, D). We performed GO analyses on significantly (p < 0.05) upregulated (1080 DEGs in females and 278 in males) or downregulated (784 DEGs in females and 227 in males) genes and found enriched pathways relevant to cardiac stress, including regulation of innate immune response and regulation of cytokine production (Fig. S3C-F).

Next, we performed integrated pathway analysis on female metabolomes and transcriptomes and, like YG8-800 male mice, found a highly significant enrichment of the ribosome pathway (Fig. S3G). After removal of ribosomal genes from the analysis, we could better detect the enrichment of proteotoxic stress pathways (autophagy, proteasome, ubiquitin mediated proteolysis), oxidative phosphorylation, and regulation of actin cytoskeleton (Fig 2E). These pathways, as well as PI3K-Akt signaling and vascular smooth muscle contraction, were also enriched in male KIKO-700 hearts (Fig. 2F). Collectively, these results suggest that approximately 50% reduction in FXN in the KIKO-700 heart is associated with cardiac stress, and metabolic and transcriptional changes consistent with ISR^mt^.

### Severe FXN deficiency in Fxn^G127V^ hearts is associated with transcriptional and metabolic alterations suggestive of cardiomyopathy and ISR^mt^

The third mouse model of FA we analyzed was a homozygous Fxn^G127V^ KI, which harbors a missense mutation in exon 4 of both *Fxn* alleles resulting in a G127V amino acid change (Fil et al., 2020). This mutation is equivalent to the human pathogenic G130V found in a small subset of FA patients (Bidichandani et al., 1997; Galea et al., 2016). Mouse embryonic fibroblasts isolated from Fxn^G127V^ mice showed severe reduction in FXN levels (5% residual), decreased mitochondrial length and increased mtDNA damage (Fil et al., 2020). Therefore, we studied the heart of this mouse as a model of severe but partial FXN loss. By western blot of whole tissue lysates (25 μg of total protein), FXN was undetectable in Fxn^G127V^ hearts (Fig 3A). However, by loading 100 μg of heart lysates, we were able to detect approximately 1% residual FXN (Fig. 3A, S2B).

**Figure 3.**
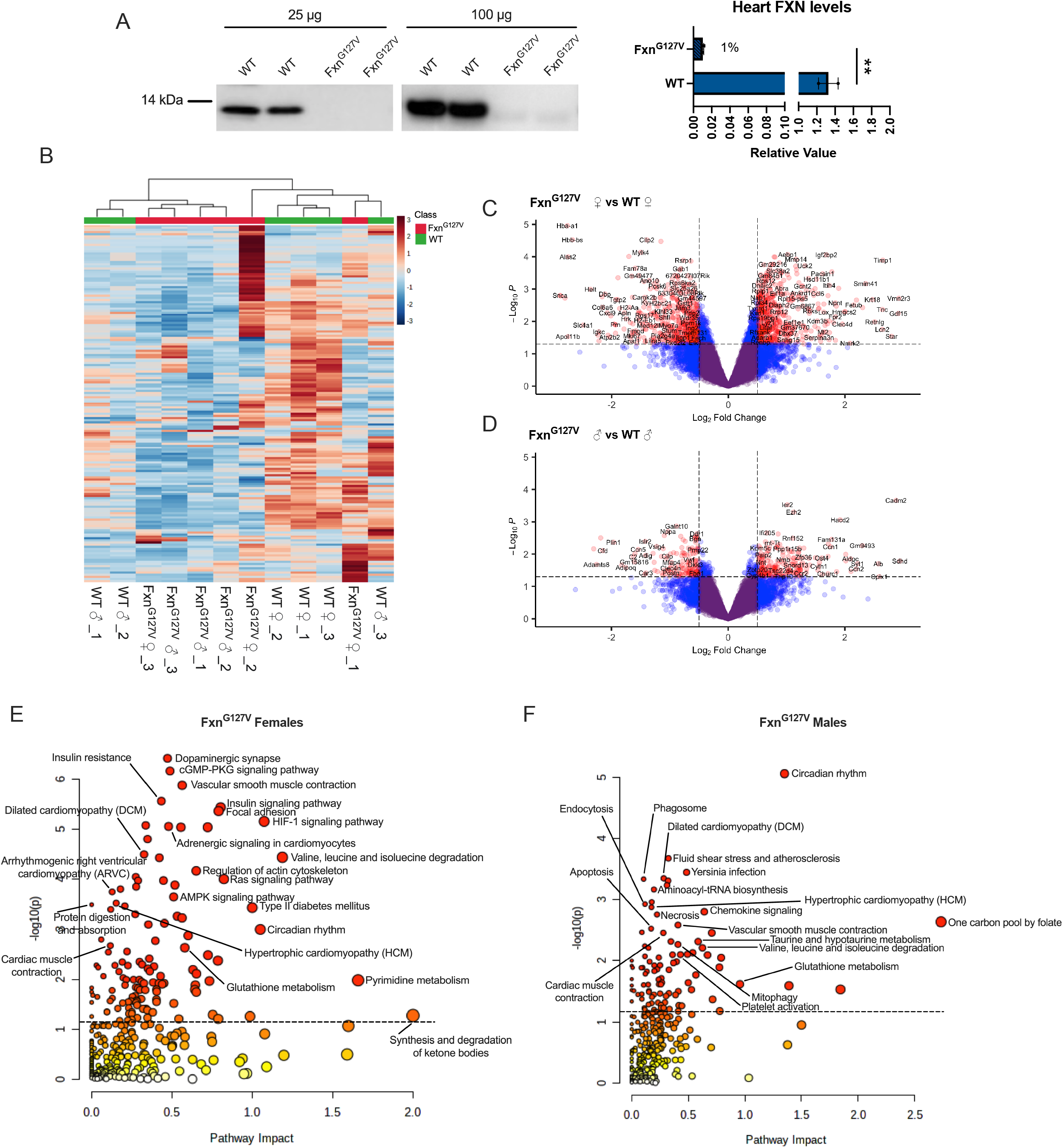
Enrichment of pathways associated with ISR^mt^ and cardiac stress in Fxn^G127V^. (A) Western blot analyses of FXN expression in WT and Fxn^G127V^ male hearts loading either 25 μg or 100 μg, and the 100 μg blot quantified to the right (n = 2) (statistical analysis was performed with unpaired two-tailed *t*-test, ** = p < 0.007, data represented as mean ± SEM). (B) Heatmap of all measured metabolites in WT Fxn^G127V^ hearts (n = 3/sex/genotype 18-months old mice). Red indicates increased abundance and blue indicates reduced abundance. (C, D) Volcano plot of gene expression in KIKO-700 females (C) and males (D) (n = 3/sex/genotype 18-months old mice). Genes in red indicate p > 0.05, FC > 0.5. (E, F) Integrated metabolomics and transcriptomics data from Fxn^G127V^ females (E) and males (F).

We performed metabolomics and transcriptomics analyses on both male and female hearts from Fxn^G127V^ and WT mice at 18 months of age. Hierarchical clustering analysis of Fxn^G127V^ and WT heart metabolites showed clustering by genotype (Fig. 3B). In both male and female Fxn^G127V^ hearts, all differential metabolites were decreased in abundance. Unbiased metabolic KEGG pathway analysis of the female Fxn^G127V^ metabolome found enrichment of metabolites involved in purine, glutathione, and amino acid metabolism (valine, leucine, isoleucine, arginine) (Fig. S4A). In male Fxn^G127V^ hearts, enriched metabolic pathways included glutathione metabolism, pantothenate and CoA biosynthesis, pyrimidine metabolism, as well as D-glutamine and D-glutamate metabolism (Fig. S4B). Transcriptomics revealed many DEGs in both female (1678 DEGs) and male (1089 DEGs) Fxn^G127V^ hearts, with an overrepresentation in females (Fig. 3C, D). GO analyses on significantly upregulated (856 in females and 514 in males) or downregulated (822 in females and 575 in males) genes showed enriched pathways known to be associated with ISR^mt^ and cardiac stress, including for example response to stress in female (Fig. S4C, D) and immune system process and tetrahydrofolate metabolic process in male Fxn^G127V^ hearts (Fig. S4E,F).

Integrated pathway analysis of significantly (p < 0.05) altered metabolites and transcripts in female Fxn^G127V^ hearts revealed significant enrichment of the ribosome pathway (Fig. S4G). After removing ribosomal genes from the analysis, several pathways associated with cardiac stress could be visualized, such as dilated cardiomyopathy, hypertrophic cardiomyopathy, vascular smooth muscle contraction, cardiac muscle contraction, and regulation of actin cytoskeleton (Fig. 3E). There were also several enriched pathways associated with ISR^mt^, including one carbon pool by folate, glutathione metabolism, mitophagy, amino acid metabolism, pyrimidine metabolism, and chemokine signaling. Several of these pathways were also enriched in integrated pathway analysis of Fxn^G127V^ male hearts, including hypertrophic cardiomyopathy, one carbon pool by folate, and glutathione metabolism (Fig. 3F). Together, these results suggest that the severe reduction in FXN expression in the Fxn^G127V^ heart leads to metabolic and transcriptional alterations associated with hypertrophic cardiomyopathy and ISR^mt^.

### Targeted transcriptomics analyses reveal markers of hypertrophic cardiomyopathy only in Fxn^G127V^ hearts

Based on the unbiased metabolomics and transcriptomics data, the three mouse models of partial FXN deficiency all shared features of cardiac stress, while pathways consistent with ISR^mt^ were only identified in KIKO-700 and Fxn^G127V^ hearts. To better understand the cardiac involvement in these models and their relationship to the human disease, we first performed a targeted analysis of the genes included in the KEGG “hypertrophic cardiomyopathy (HCM)” pathway (pathway mmu05410). This pathway encompasses a total of 91 genes, 70 of which were detected by RNAseq in most groups (except YG8-800 mice, with only 59 genes detected). Approximately 20% of HCM pathway genes were significantly different (p < 0.05) in Fxn^G127V^ female hearts relative to WT mice, 14% in Fxn^G127V^ males, 12% in KIKO-700 females, 5% in KIKO-700 males, and 14% in YG8-800 males (Fig. 4A). Surprisingly, there was little overlap among groups and none of these DEGs were common to all groups. Therefore, to investigate the putative gene expression changes more specifically associated with FA cardiomyopathy, we analyzed a subset of 15 genes, which were found to be altered in human induced pluripotent stem cells-derived cardiomyocytes from FA patients (Li et al., 2019) or in a mouse model of heart-specific FXN cKO (Perdomini et al., 2014). Only in the Fxn^G127V^ female hearts we found increased expression of natriuretic peptide precursor b (*Nppb*), a fetal gene that is overexpressed under cardiac stress conditions (Fig. 4B). In addition to this gene list, only in Fxn^G127V^ female hearts, we found upregulation of reticulon 4 (*Rtn4*), encoding an ER protein involved in sphingolipid regulation (Sasset et al., 2022), which is known to be elevated in mouse models of hypertrophic cardiomyopathy (Sasagawa et al., 2016). KIKO-700 males showed a downregulation of natriuretic peptide precursor a (*Nppa*) and myosin heavy chain beta (*Myh7*) (Fig. 4B), which was not consistent with development of cardiomyopathy. Hearts from all other mouse groups did not display consistent changes in the transcriptional profiles of this subset of genes. Overall, these results suggest a sexual dimorphism in the expression of cardiomyopathy markers in Fxn^G127V^ mice, while both KIKO-700 and YG8-800 did not display alterations in the expression of the subset of FA-related hypertrophic cardiomyopathy genes.

**Figure 4.**
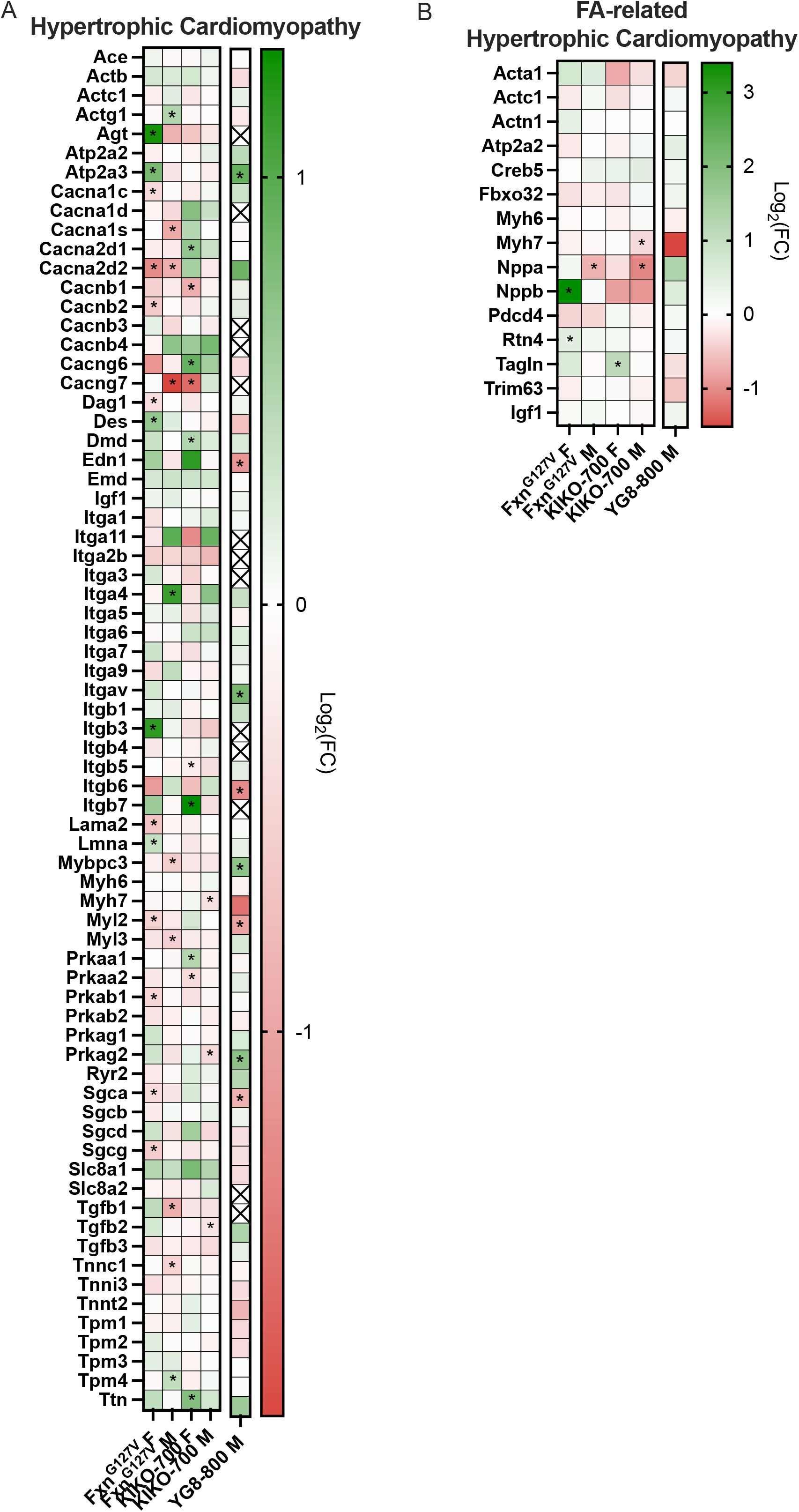
Hypertrophic cardiomyopathy marker expression only in Fxn^G127V^ hearts. (A) Targeted gene expression analysis of the KEGG pathway (mmu05410) “Hypertrophic Cardiomyopathy” represented in a heatmap where green indicates increased gene expression and red indicates decreased and * = p < 0.05. An “X” indicates that a gene was not detected by RNAseq. (B) Heatmap showing targeted gene expression of genes associated with FA-related hypertrophic cardiomyopathy. In all heatmaps, expression is represented as Log_2_(fold change [FC]). n = 3/sex/genotype 18-months old FXN^G127V^ and KIKO-800 and n = 4 18-months old YG8-800.

The loss of FXN leads to iron dysregulation in the heart, including increased iron uptake, iron accumulation in mitochondria, and ferroptosis (Martelli and Puccio, 2014). To understand the effects of partial FXN deficiency on heart iron metabolism, which could underlie cardiomyopathy, we performed targeted gene expression analyses of genes related to iron, including those involved in transferrin-dependent and -independent iron uptake, iron storage and export, and mitochondrial iron import. Only in Fxn^G127V^ female hearts there was an increase in transferrin receptor (*Tfrc*), ferritin (*Fth1*), and ferroxidase (*Cp*) gene expression relative to WT (Fig. S5A). Furthermore, among genes involved in Fe-S cluster biogenesis and heme metabolism, we found decreased expression of *Nfs1, Nubpl*, and *Fech* (Fig. S5B, C) only in Fxn^G127V^ female hearts. In males, Fxn^G127V^ and KIKO-700 hearts had increased expression of aminolevulinic acid synthase 1 (*Alas1*), the heme biosynthesis rate limiting mitochondrial enzyme, while YG8-800 males had increased expression of heme oxygenase-1 (*Hmox1*), which catalyzes the first step in heme degradation (Fig. S5C). Overall, this data indicates that only a few genes of iron metabolism are affected in a genotype- and sex-dependent manner and that these changes are more prominent in Fxn^G127V^ female hearts.

The adult heart relies primarily on β-oxidation of fatty acids to provide substrates for mitochondrial OXPHOS. Under stress, the heart alters metabolic fuel sources to support contraction shifting its energy metabolism towards glycolysis (Lopaschuk and Jaswal, 2010; Lopaschuk et al., 2010). The integrated pathway analyses performed on metabolites and transcripts of the heart of the three mouse models of partial FXN deficiency suggested the involvement of several pathways related to energy metabolism (Figs. 1F, 2E, F, 3E, F). Therefore, to better define the potential link between cardiomyopathy and energy metabolism, we performed a targeted analysis of the heart transcriptome and metabolome in the three models. We evaluated the expression of genes of the KEGG “β-oxidation” pathway (module M00087). This pathway encompasses a total of 13 genes, 11 of which were detected by RNA sequencing (RNAseq) in most groups (except YG8-800 mice, with only 10 genes detected). Majority of the genes were not differentially expression, however *Hadh* was downregulated in FXN^G127V^ female hearts and *Acaa1a* was upregulated in KIKO-700 female hearts (Fig. 5A). Furthermore, the levels of β-oxidation intermediates were mostly unchanged, except for a significant decrease of L-acetylcarnitine in Fxn^G127V^ and KIKO-700 females (Fig. 5B). We also looked at the expression of genes of the KEGG “glycolysis” pathway (module M00001). This pathway encompasses a total of 28 genes, 20 of which were detected by RNAseq in most gropus (except YG8-800 mice, with only 18 genes detected). Fxn^G127V^ females showed a significant increase of the expression of *Pfkp* (phosphofructokinase, platelet) (Fig. 5B). Notably, *Pfkp* overexpression was previously described in a model of hypertrophic cardiomyopathy induced by pressure overload (Vigil-Garcia et al., 2021). In addition, we detected decreased expression of *Hk2* (hexokinase 2) and *Eno3* (enolase 3) in Fxn^G127V^ females. In these mice, we found a significant reduction in G3P (glyceraldehyde 3-phosphate) levels (Fig. 5D), which were also decreased in Fxn^G127V^ males and KIKO-700 females. Fxn^G127V^ male hearts also showed a significant reduction in the level of pyruvate. KIKO-700 females had reduced levels of 3PG (3-phosphoglyceric acid) and PEP (phosphoenolpyruvic acid). Overall, these findings indicate that metabolic consequences of cardiac stress were manifested in several groups, but mostly in Fxn^G127V^ female hearts.

**Figure 5.**
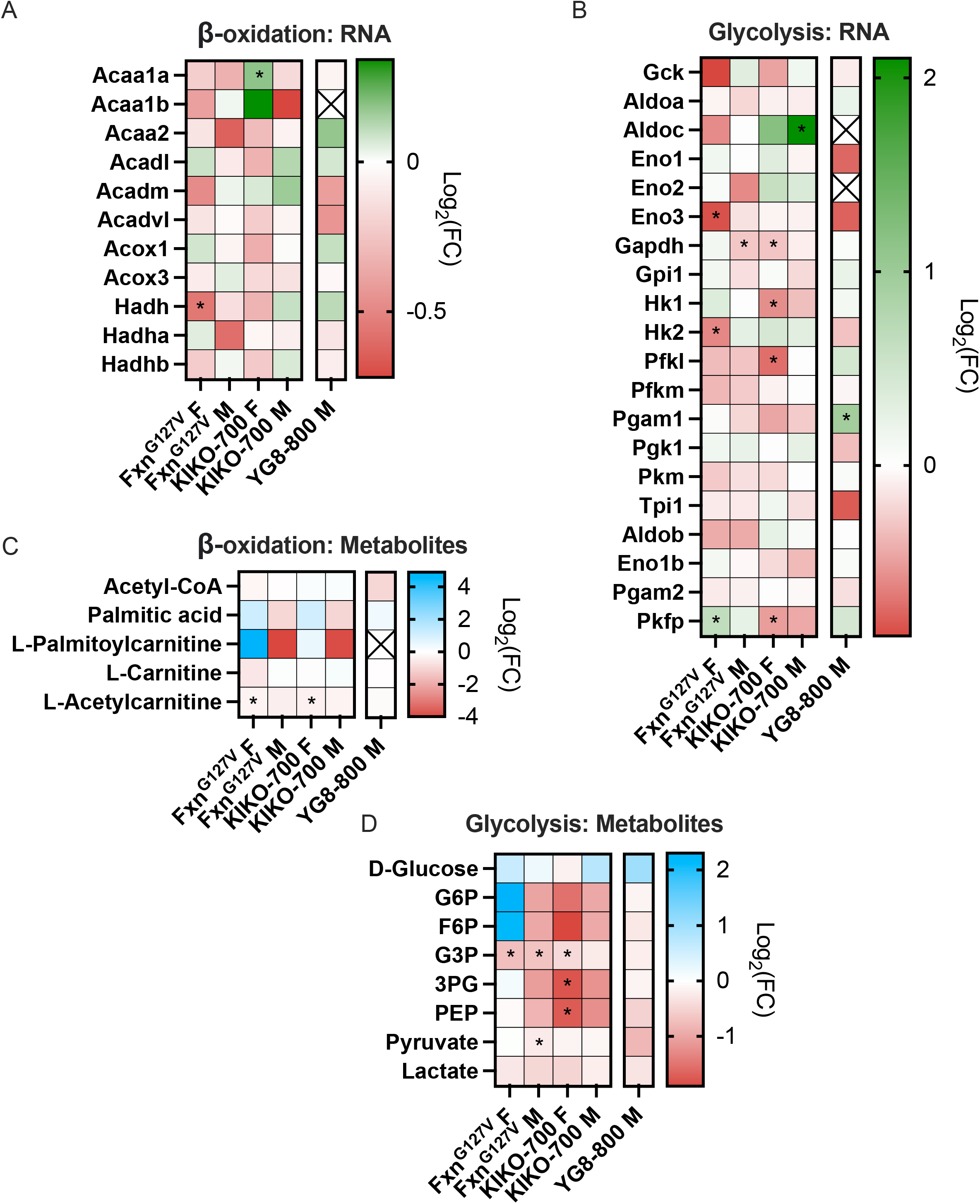
Normal expression of β-oxidation and glycolysis genes and metabolites in all three FXN-deficiency models. (A, B) Heatmap showing the expression of genes related to (A) β-oxidation (KEGG module M00087) and (B) glycolysis (KEGG module M00001). (C, D) Heatmap showing the abundance of metabolites related to (C) β-oxidation and (D) glycolysis, where blue indicates increased abundance and red indicates decreased abundance. Abbreviations: G6P, glucose-6 phosphate; F6P, fructose-6 phosphate; G3P, glycerol-3-phosphate; 3PG, 3-phosphoglyceric acid; PEP, phosphoenolpyruvic acid. n = 3/sex/genotype 18-months old FXN^G127V^and KIKO-800 and n = 4 18-months old YG8-800. * = p < 0.05.

### Transcriptional and metabolic profiles of ISR^mt^ are only evident in Fxn^G127V^ hearts

Unbiased integrated pathway analysis of the three models of FXN deficiency highlighted enrichment of pathways that may suggest ISR^mt^ activation in Fxn^G127V^ and KIKO-700 hearts. Therefore, to characterize ISR^mt^-related pathways more in depth, we first performed targeted analysis of the expression of canonical ISR^mt^-related genes (Forsstrom et al., 2019). We found upregulation of the transcription factors *Atf4* in female Fxn^G127V +^ hearts and *Atf5* in both males and females, but not in the other mouse models (Fig. 6A). Furthermore, in Fxn^G127V^ hearts there was an upregulation of the expression of cytokine *Gdf15*, transcription factor *Trib3*, and asparagine synthetase (*Asns*), indicative of a cardiac ISR^mt^ activation (Dogan et al., 2014; Forsstrom et al., 2019; Kaspar et al., 2021; Khan et al., 2017; Kuhl et al., 2017; Nikkanen et al., 2016; Sayles et al., 2022).

**Figure 6.**
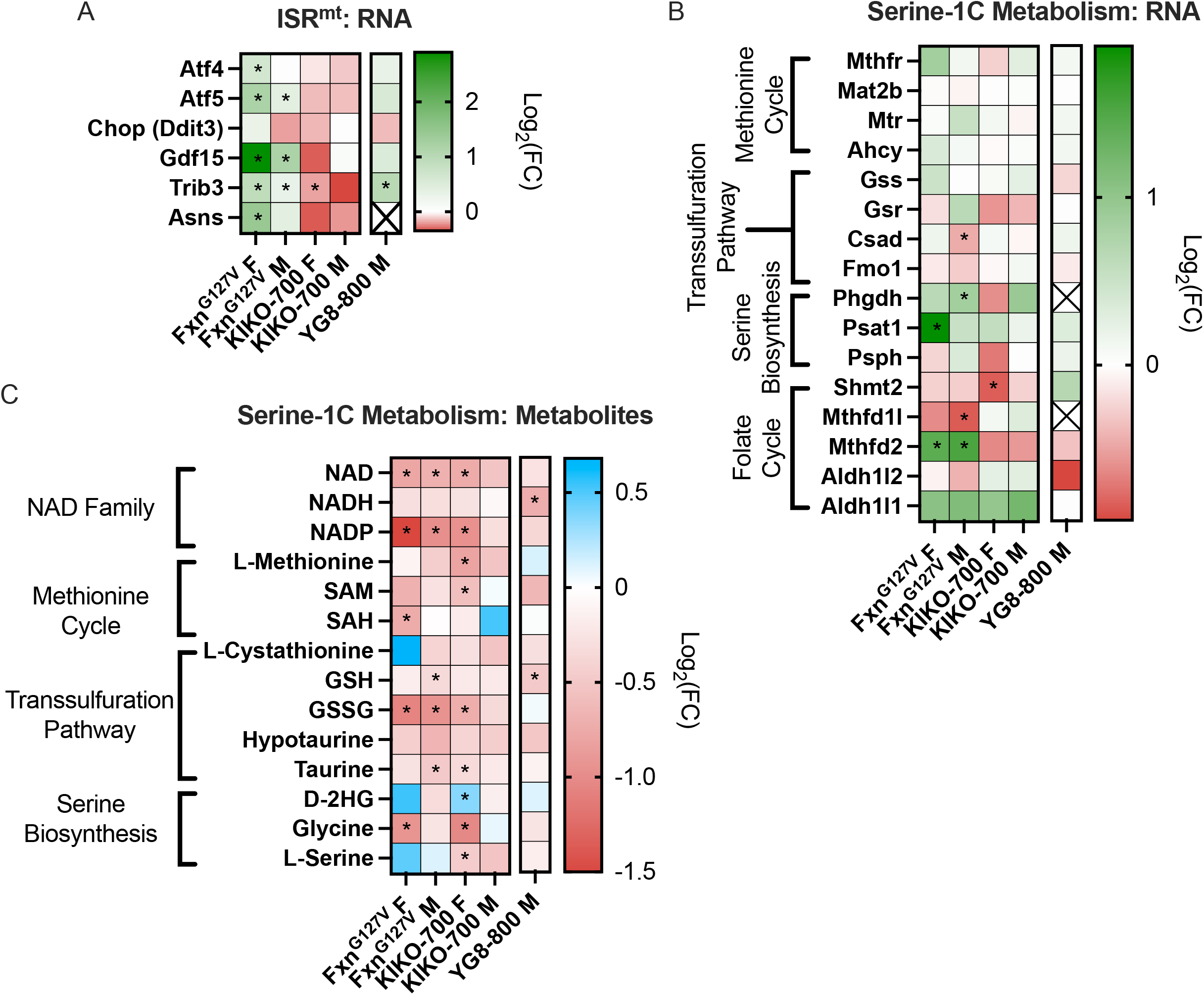
Markers of ISR^mt^ activation and 1C metabolism alterations in Fxn^G127V^ hearts. (A, B) Heatmap showing the expression of genes related to (A) ISR^mt^ and (B) serine-1C metabolism. (C) Heatmap showing the abundance of metabolites related to serine-1C metabolism, including NAD family, methionine cycle, transsulfuration pathway, and serine biosynthesis. n = 3/sex/genotype 18-months old FXN^G127V^ and KIKO-800 and n = 4 18-months old YG8-800. * = p < 0.05.

Metabolic remodeling associated with ISR^mt^ includes the upregulation of serine-1C metabolism. We examined all genes of the serine-1C metabolism and found that most of these genes were unchanged in the KIKO-700 and YG8-800 models (Fig. 6B). In Fxn^G127V^ hearts, however, there was a significant upregulation of serine biosynthesis pathway genes, *Phgdh* in males and *Psat1* in females, as well as folate cycle genes, *Mthfd2* in males and females (Fig. 6B). The elevation of serine-1C metabolism promotes the transsulfuration pathway for the methionine cycle to enhance GSH production and antioxidant defenses (Wu and Storey, 2021). However, the levels of GSH and GSSG in Fxn^G127V^ hearts were decreased (Fig. 6C), suggesting that the glutathione biosynthetic pathway is unable to meet the demands of increased oxidative stress. Of note, NAD and NADP levels were decreased in all three models, by a greater degree in Fxn^G127V^ hearts (Fig. 6C). Although the metabolic consequences of these alterations remain to be further elucidated, NAD depletion was reported in a mouse model of cardiac-specific FXN cKO in association with perturbations of SIRT1 activity and the NAD salvage pathway (Chiang et al., 2021). Next, we analyzed the expression of all detected GSH-linked antioxidant peroxidases (*Gpx*) and S-transferases (*Gst*) and only found increased expression of *Gpx3* and *Gpx8* in Fxn^G127V^ hearts (Fig. S6A). Furthermore, we investigated the expression of other antioxidant enzymes, including peroxiredoxins (*Prdx*), superoxide dismutases (*Sod*), and Nrf2-driven genes and found no changes (Fig. S6B). In addition, we analyzed the expression of NADPH oxidase components (NOX2/4), as increased expression of these enzymes has been associated with ISR^mt^ activation and linked to reactive oxygen species production and stress signaling (Chen et al., 2012; Nabeebaccus et al., 2017; Zhao et al., 2015), and did not see significant alterations in any of the models, except for *Ncf1* (NOX2 complex subunit) upregulation in Fxn^G127V^ male hearts (Fig. S6C). Overall, increased *Atf4/5* expression and reduced GSH levels indicate moderate ISR^mt^ and oxidative stress in Fxn^G127V^ hearts.

### Early ISR^mt^ activation in Fxn^G127V^ hearts in the absence of defects in the activity of Fe-S cluster-dependent enzymes

Previous studies have shown that ISR^mt^ progresses in temporal stages (Forsstrom et al., 2019) and, when chronically activated, leads to sustained metabolic alterations that contribute to cardiomyopathy (Sayles et al., 2022). Since aged 18-months old Fxn^G127V^ mice showed the most evidence of cardiac ISR^mt^ activation among the three mouse models we studied, we wanted to investigate ISR^mt^ markers at an earlier time point in these mice. At 6-months of age, we found an increase in the expression of *Fgf21* in female and male Fxn^G127V^ hearts and an upregulation of *Gdf15* and *Mthfd2* in males (Fig. 7A). However, we did not find alterations in the expression of other established ISR^mt^ genes, including *Atf4, Atf5, Psat1*, and *Asns*. These findings are consistent with the initial disease phase of ISR^mt^, originally described in skeletal muscle (Forsstrom et al., 2019), in which elevation of a key mediator of metabolic remodeling *Fgf21*, accompanied by increased *Gdf15* and *Mthfd2* expression, was shown to be involved in the progression of ISR^mt^.

**Figure 7.**
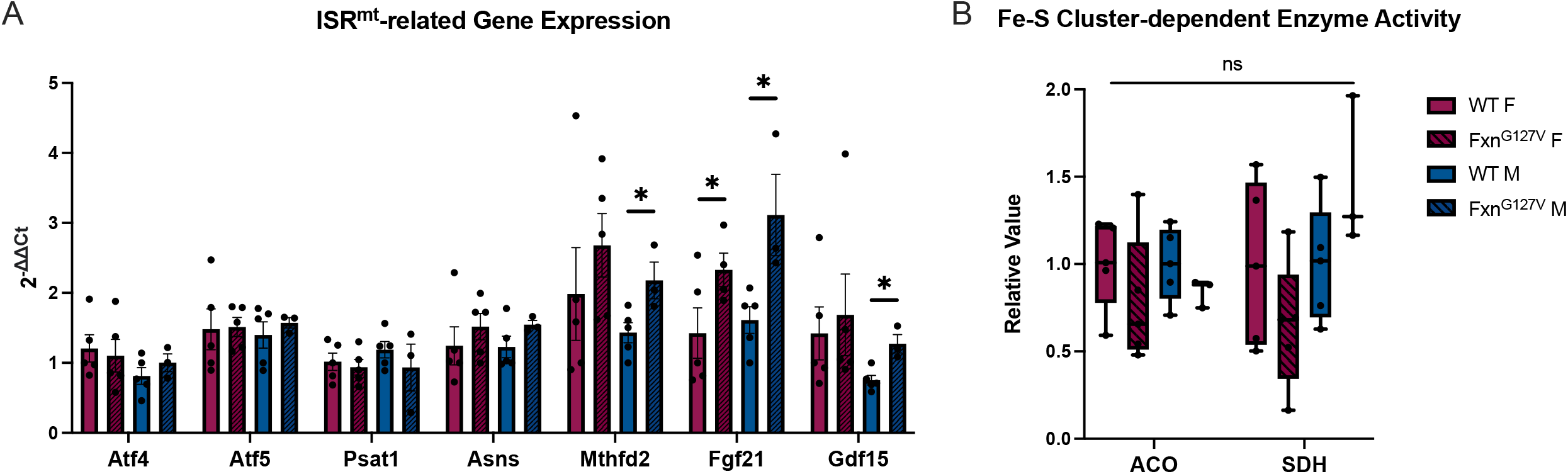
Early markers of ISR^mt^ in male Fxn^G127V^ hearts. (A) mRNA expression of genes related to ISR^mt^ (n = 5/sex 6-months old mice, except n = 3 for FXN^G127V^ males). Expression is normalized to *Gadph* expression. Data represented as mean ± SEM. (B) Activity of Fe-S cluster-dependent enzymes ACO and SDH in 6-months old mice. Values are relative to WT littermate controls. Data were represented as a box-and-whisker plot: whiskers extend from the minimum to maximum values; lines indicate median. Statistical analyses were performed with unpaired two-tailed *t*-test, * = p < 0.05.

In patients with FA, insufficient levels of FXN leads to a decreased Fe-S cluster biogenesis, resulting in an impairment of Fe-S cluster-dependent enzyme activity, including aconitase (ACO) and succinate dehydrogenase (SDH) (Puccio and Koenig, 2000; Rotig et al., 1997). Therefore, we wanted to investigate if the severe FXN deficiency seen in Fxn^G127V^ hearts (<1% residual FXN, Fig. 3A) causes an enzymatic defect that may contribute to mitochondrial dysfunction and ISR^mt^. Surprisingly however, both ACO and SDH activity were not decreased in Fxn^G127V^ hearts compared to sex and age matched controls at 6 months (Fig. 7B). Together, this data suggests that very low levels of FXN are sufficient to maintain Fe-S-dependent enzymatic function for at least 6 months and that ISR^mt^ can be initiated independently of the function of these enzymes.

## Discussion

The ISR^mt^ is an evolutionarily conserved adaptive response designed to help the heart face periods of acute stress (Eckl et al., 2021). However, if chronic and unresolved, as in the case of genetically determined mitochondrial diseases, it can become “maladaptive”, and cause detrimental metabolic imbalances that could contribute to heart failure (Smyrnias, 2021). One-carbon metabolism is a major metabolic pathway upregulated under ISR^mt^ conditions (Mehrmohamadi et al., 2014; Nikkanen et al., 2016). Interestingly, metabolic perturbations that included markers of dysregulated 1C metabolism were found in serum from FA patients (O’Connell et al., 2022). Although the cardiac origin of these markers remains to be established, these findings suggest the involvement of ISR^mt^ in FA. Furthermore, ISR^mt^ activation has been reported in models of genetic deletion or silencing of cardiac FXN (Huang et al., 2013; Tong et al., 2022; Vasquez-Trincado et al., 2021). While these models suggest that chronic ISR^mt^ is associated with FXN loss, the complete loss of FXN does not accurately reflect the human disease, where there are variable amounts of residual FXN in the heart. Therefore, investigation of ISR^mt^ and a clearer understanding of its metabolic consequences in mouse models of FA that are genetically more representative of the condition in human patients is needed, to determine if this pathway could be targeted therapeutically. Here, we investigated cardiac ISR^mt^ in YG8-800, KIKO-700 and Fxn^G127V^ mouse models, which have partial FXN deficiency.

The three models had varying degrees of cardiac FXN deficiency, with the most severe reduction observed in Fxn^G127V^ hearts, in which there was approximately 1% residual FXN. Interestingly, despite having longer GAA expansions, the levels of FXN in both YG8-800 and KIKO-700 hearts were comparable to the FXN levels reported in the YG8sR (200 GAA repeats) (Anjomani Virmouni et al., 2015) and KIKO-230 (230 GAA repeats) (Miranda et al., 2002) mouse models. This suggests that mouse models of GAA expansion do not fully recapitulate the human disease, in which there is an inverse correlation between expansion size and levels of FXN (Koeppen, 2011).

Unbiased analyses of the transcriptomes and metabolomes of hearts from these FA mouse models identified oxidative phosphorylation and fatty acid metabolism among the most significantly altered pathways in YG8-800, while 1C and amino acid metabolism were among the most altered pathways in KIKO-700 and Fxn^G127V^. Pyrimidine and glutathione biosynthesis were also altered in Fxn^G127V^ hearts. Furthermore, we observed an enrichment of pathways related to cardiac stress and cardiomyopathy in all three models. These findings provide evidence of cardiac ISR^mt^-related perturbations in these mice, although overt symptomatic cardiomyopathy has not been described in these mouse models. Since integrated pathway analysis of gene expression and metabolite abundance suggested cardiac ISR^mt^, we performed targeted analyses of known mRNA markers of cardiomyopathy, such as *Myh6/7* and *Nppa/b*. However, only the Fxn^G127V^ hearts showed an elevation of some cardiomyopathy markers, with prevalence in females. Similarly, mRNA markers of ISR^mt^, such as *Atf4/5* and *Mthfd2*, were only upregulated in Fxn^G127V^ hearts. The most evident difference between the Fxn^G127V^ and the YG8-800 and KIKO-700 mouse models is the level of residual FXN. It was suggested, based on findings in the inducible *Fxn* silencing model, that 20% residual FXN is sufficient to prevent OXPHOS impairment, cardiac dysfunction, and related metabolic responses (Vasquez-Trincado et al., 2021). With the caveat that this model is a post-natal silencing of *Fxn*, unlike the human disease where presumably patients have reduced FXN levels since embryonic development, this data suggests the threshold for cardiac involvement of ISR^mt^ is below 20% residual FXN in the mouse. Both the YG8-800 and KIKO-700 have more than 20% FXN in the heart, likely explaining the lack of alterations of specific cardiac stress markers.

Hearts from the Fxn^G127V^ mice displayed markers of ISR^mt^ progression. Based on a time course analysis of ISR^mt^ in skeletal muscle of a mouse model of mitochondrial myopathy, early and late stages were described (Forsstrom et al., 2019). In the 6-month-old cohort of Fxn^G127V^ male mice, we found increased expression of markers of early ISR^mt^, including *Mthfd2, Gdf15*, and *Fgf21*. These markers were also shown to be upregulated prior to *Atf4/5* gene expression changes in a mouse model of mitochondrial cardiomyopathy (Sayles et al., 2022). Despite the extremely low levels of FXN in Fxn^G127V^ hearts, the activities of Fe-S cluster-dependent enzymes ACO and SDH were unchanged at 6-months of age, suggesting that ISR^mt^ in the mouse heart can start prior to the onset of bioenergetic defects, like a previously described model of mitochondrial cardiomyopathy (Sayles et al., 2022). At 18-months of age, both male and female Fxn^G127V^ hearts express markers of ISR^mt^. The earlier activation of ISR^mt^ in males suggests sexual dimorphism, due to yet unknown mechanisms. Future studies will investigate the time of onset of ISR^mt^ in the female Fxn^G127V^ hearts and the potential role of sex hormones in delaying ISR^mt^ activation.

Our findings highlight important differences among the Fxn^G127V^ and the *Fxn* GAA repeat expansion models (YG8-800 and KIKO-700). We propose that only the Fxn^G127V^ mutation causes sufficiently low levels of FXN in the mouse heart to trigger the upregulation of specific ISR^mt^ genes. The dramatic decrease (<1% residual FXN), however, did not result in a fatal cardiomyopathy, nor did it cause ACO and SDH enzymatic defects. Therefore, the mechanisms of ISR^mt^ induction in response to FXN loss in this model remain to be elucidated. A putative mechanism could involve defective FXN maturation due to the Fxn^G127V^ mutation, which could be a source of proteotoxic stress in mitochondria.

In summary, we showed that ISR^mt^ may arise in the heart of mouse models of FA that recapitulate the human disease, which is associated with partial FXN deficiency rather than complete genetic depletion. However, there are limitations to these models, as they only develop a partial cardiac mitochondrial stress response, which complicate studies of mechanism and clinical implications. These findings further highlight the difficulty in modeling FA cardiomyopathy in murine models, as the threshold for preservation of cardiac function is likely as low as 1% residual FXN.

## Methods

### Mouse models

All animal procedures were conducted in accordance with Weill Cornell Medicine, University of Alabama at Birmingham and IGBMC Animal Care and Use Committees and were performed according to the Guidelines for the Care and Use of Laboratory Animals of the National Institutes of Health. YG8-800 mice were generated from the previously published YG8sR (Anjomani Virmouni et al., 2015) and are available from The Jackson Laboratory (Fxn^em2.1Lutzy^ Tg(FXN)YG8Pook/800J, Stock# 030395). CRISPR/Cas9-generated Fxn^G127V^ mice were previously generated (Fil et al., 2020). Mice were euthanized by cervical dislocation.

The KIKO-700 mouse strain was generated using a standard homologous recombination approach on the C57BL/6 background. First, a 12.3 kbp genomic DNA used to construct the targeting vector was subcloned from a positively identified C57BL/6 fosmid clone (WI1-1938E18). The region was designed such that the 5’ homology arm extends ~7.9 kbp to the human sequence with GAA repeats and the FRT-flanked neomycin (Neo) cassette. The 3’ homology arm extends about 3.4 kbp from the Neo cassette. The human sequence with GAA repeats and Neo cassette was inserted 1,400 bp downstream of mouse *Fxn* exon 1. The human sequence containing ~ 700 GAA repeats was amplified by PCR using genomic DNA isolated from FRDA patient cells as described (Li et al., 2016). The targeting vector was confirmed by restriction analysis and sequencing after each modification step. Subsequently, the construct containing ~700 GAAs was transfected into FLP C57BL/6 embryonic stem cells (ES) (Ingenious) and positive clones were identified by neomycin selection. Genomic DNA was isolated from 400 ES clones and screened for correct integration and appropriate length of the GAA tract. Only 7% of clones harbored a tract of ~700 GAAs. The selection cassette was removed using FLP recombinase. Five separate injection sets were conducted resulting in successful generation of two chimeras. Subsequently, chimeras were bred with WT B6 mice to obtain fully heterozygous KI (*Fxn^+/700^*) animals. The FLP allele was removed by breeding with wild type C57BL/6 animals. Finally, KI *Fxn^+/700^* mice were crossed with heterozygous *Fxn* mice exon 4 deleted (*Fxn^+/-^* (B6.129(Cg)-Fxntm1Mkn/J, The Jackson Laboratory stock no. 016842) to obtain *Fxn^700/-^* (KIKO-700) study animals. All animals included in this study were genotyped and length of the GAA repeats was verified by repeat PCR. No significant germline instability was observed. Detailed characterization of neurobehavioral phenotypes of the *Fxn^700/-^* (KIKO-700) mice will be described in Li, *et al*. (manuscript in preparation).

### RNA Sequencing

RNA was extracted from heart tissue using TRIzol (Life Technology) and the RNeasy Mini Kit (Qiagen), according to manufacturer’s instructions. 3’RNAseq libraries were prepared from 500 ng of RNA per sample using the Lexogen QuantSeq 3’ mRNA-Seq Library Prep Kit FWD for Illumina and pooled for reduced run variability. Libraries were sequenced with single-end 86 bps on an Illumina NextSeq500 sequencer (Cornell Genomics Facility). All computations were performed in the R statistical environment (version >4.2.0; https://www.R-project.org/). Raw sequence reads were processed using ShortRead package (version 1.54.0) (Morgan et al., 2009). Trimmed reads were aligned to the mouse genome assembly GRCm38.94 using the Rsubread package (version 2.10.5) (Liao et al., 2019) with default parameter settings. The Rsubread::featureCounts function was used to assign mapped stranded sequencing reads to genes and count features. The Limma package (version 3.52.1) (Ritchie et al., 2015) was used to obtain normalized and variance stabilized counts and to calculate differential gene expression. Pathway analysis for all gene expression data was performed with the gprofiler2 (Kolberg et al., 2020) and clusterProfiler packages (Wu et al., 2021), using the GO Biological Process and KEGG databases. A false discovery rate corrected p value of < 0.05 was used to determine significance. Pathways shown in the figures were condensed using the simplify function from the clusterProfiler package (Wu et al., 2021) to merge terms with more than 40% overlapping annotated genes.

### Metabolomics

Untargeted metabolomics of heart tissue was performed at Weill Cornell Medicine Meyer Cancer Center Proteomics & Metabolomics Core Facility. Briefly, 15 mg of cardiac tissue was homogenized in 80% methanol (Sigma) using Tissue Tearer (BioSpec) on dry ice. Samples were incubated at −80°C for 4 hours. Homogenates were then centrifuged at 14,000 rcf for 20 min at 4°C. The supernatant was extracted and stored at −80°C for mass spectrometry. Analysis of metabolite changes was performed with MetaboAnalyst (version 5.0) (Xia et al., 2009), which included the following: fold change analyses, heat map, pathway enrichment analysis and integrated pathway impact analysis. Integrated pathway impact analysis was performed with hypergeometric test and with degree centrality topology applied.

### ELISA

Human or mouse FXN levels were measured by ELISA (for human: Abcam; ab176112 and or mouse: Abcam; ab199078) according to manufacturer’s instructions. Briefly, protein was extracted from homogenized heart tissue using 1X Cell Extraction Buffer PTR. Standards or protein lysates were incubated with the Antibody Cocktail for one hour at room temperature on a plate shaker set to 400 rpm. Following washing and the addition of TMB Development Solution and Stop Solution, absorbance was recorded with an OD at 450 nm using the PowerWave XS (Biotek). Human and mouse FXN levels were normalized to their respective standard curve measurements.

### Western Blotting

Protein concentration was determined by the Bradford protein assay (Bio-Rad). Total heart lysates (25, 75 or 100 μg) were denatured in 1X Laemmli Buffer (Bio-Rad) containing 2-Mercaptoethanol (Sigma) at 95°C for 10 min and separated by electrophoresis in a 4–12% SDS–PAGE gel (Bio-Rad) and transferred to a PVDF membrane (Bio-Rad). Blots were incubated in 3% BSA in TBS with 1% Tween-20 for one hour at room temperature. Primary antibodies were incubated overnight at 4°C. Secondary antibodies were incubated for 45 min at room temperature. In all blots, proteins were detected using Clarity Western ECL Blotting Substrates (Bio-Rad) and imaged on ChemiDoc Touch (Bio-Rad). Normalization of FXN levels were determined by Poncaeu S staining (Sigma). The following antibodies were used: monoclonal mouse-anti-frataxin (1:1000; Milipore Sigma; clone 4F9; MABN2313) and polyclonal rabbit-anti-frataxin (1:1000; ProteinTech; 14147-1-AP).

### Quantitative PCR

RNA was extracted from heart tissue using TriZol (Life Technology), according to manufacturer’s instructions. Total mRNA (10 μg) was used for reverse transcription with SuperScript IV Reverse Transcriptase (Thermo Fisher Scientific) in 100 uM DTT, 50 μM oligo dT, 10 mM dNTP, and 40 U/μl RNAsin in a total volume of 10 μl. A PCR was performed to amplify exons. Primer sequences can be found in Table 1. Quantification of the RT-PCR product was obtained on a LightCycler 480 (Roche).

**Table 1.**
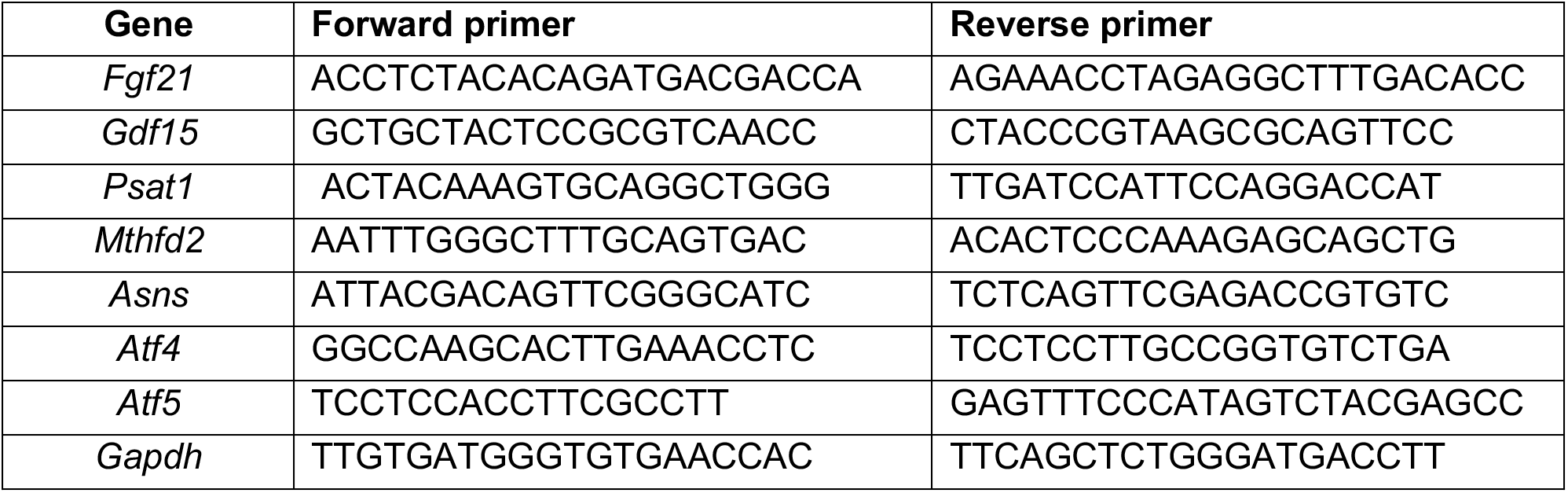
qPCR primer sequences.

### Enzymatic activity assays

ACO and SDH activity were measured as previously described (Puccio et al., 2001). Briefly, proteins were isolated from 10 mg heart tissue in 50 μl extraction buffer (10 mM KH2PO4, 2 mM EDTA, 1 mg/ml BSA). ACO activity was measured at 240 nm following the addition of 5 mM citrate. SDH activity was measured at 600 nm following the addition of 200 mM ATP, 320 mM KCN, 5 mM succinate, 0.05 mM decylubiquinone. SDH activity was normalized to isocitrate dehydrogenase activity. All reagents were from Sigma, except for KCN (Prolabo). Activity was measured on a Cary 50 Scan UV Visible Spectrophotometer (Varian).

### Data analysis

The number of animals (biological replicates) for transcriptomics and metabolomics performed at 18 months was 3/sex/genotype for FXN^G127V^ and KIKO-700 and 4/genotype for YG8-800. Biological replicates for enzymatic assays and qPCR were 5/sex/genotype (except FXN^G127V^ males, which were n = 3) at 6 months. Statistical analyses were performed using GraphPad Prism (GraphPad Software version 9.1.1). Two-group comparisons were analyzed by the two-tailed *t-*test. Differences were considered statistically significant with a p value < 0.05. Data are expressed as the mean ± SEM. Pathway analysis was performed using MetaboAnalyst using the hypergeometric test and relative-betweenness centrality. Joint pathway analysis was performed with hypergeometric test with degree centrality topology applied and these data integrated based on pathway level combined p values for all pathways.

## Acknowledgments

We acknowledge the Weill Cornell Medicine Meyer Cancer Center Proteomics & Metabolomics Core Facility and the Cornell Genomics Facility.

## Data availability

Transcriptomics data is available through Gene Expression Omnibus gene repository with the dataset identifier *in progress*. Metabolomics data was deposited to the Metabolomics Workbench with the study ID *in progress*.This paper does not report original code. Any additional information required to reanalyze the data reported in this paper is available from the lead contact upon request.

## Author contributions statement

N.M.S. performed experiments, analyzed results, and participated in experimental design and manuscript preparation; J.S.N. performed experiments, analyzed results, and participated in experimental design and manuscript preparation; J.A. performed experiments; N.D. performed experiments; J.L. performed experiments; M.N. participated in experimental design, analyzed results and manuscript preparation; H.P. participated in experimental design and manuscript preparation; G.M. participated in experimental design and manuscript preparation.

---

**Figure S1.**
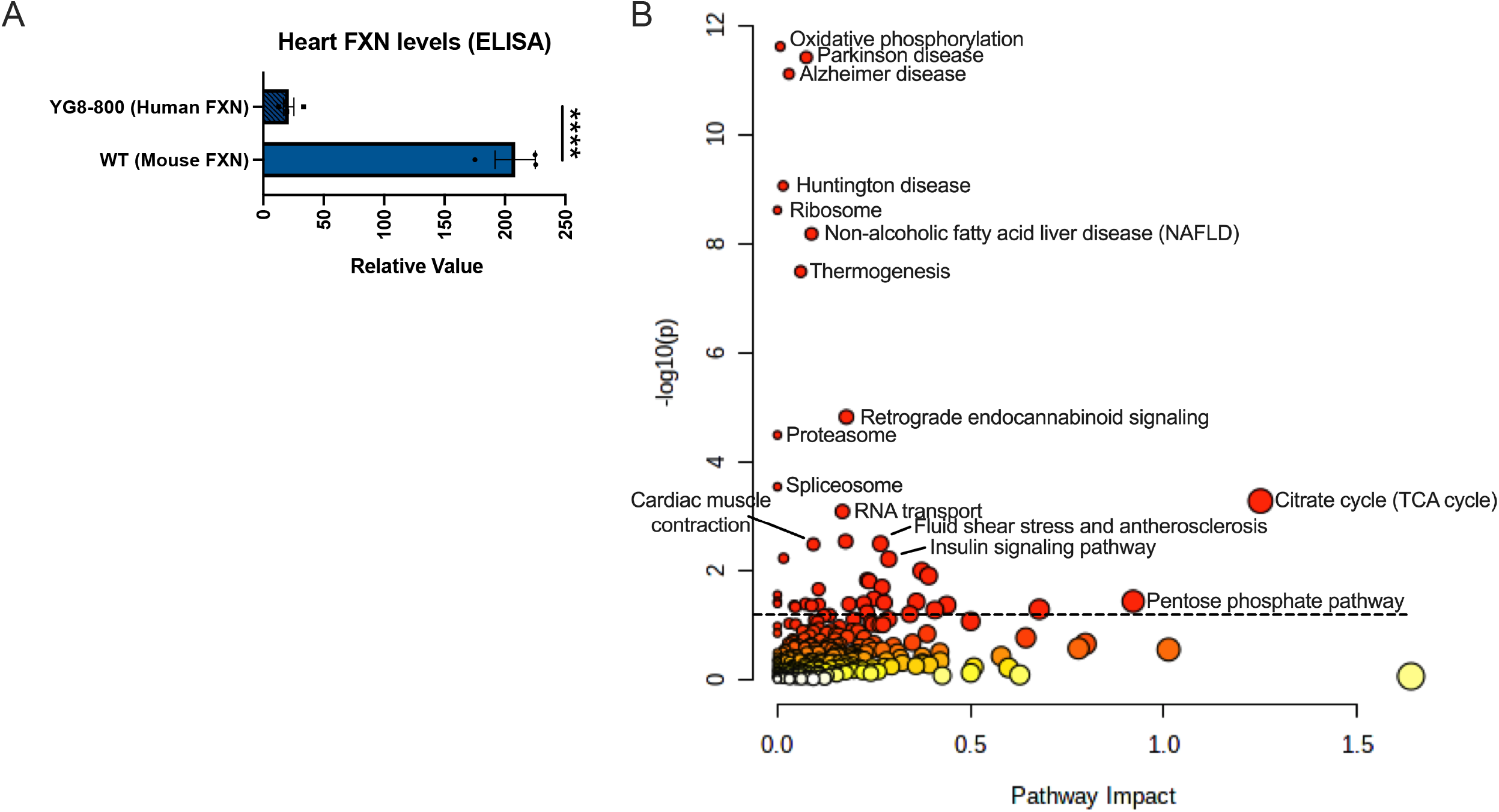
Ribosomal pathway enrichment in YG8-800 joint-pathway analysis. (A) Measurement of mouse and human FXN by ELISA (statistical analysis was performed with unpaired two-tailed *t*-test, **** = p < 0.0001, data represented as mean ± SEM). (B) Integrated metabolomics and transcriptomics data from YG8-800 males including ribosomal genes. n = 4/genotype 18-months old.

**Figure S2.**
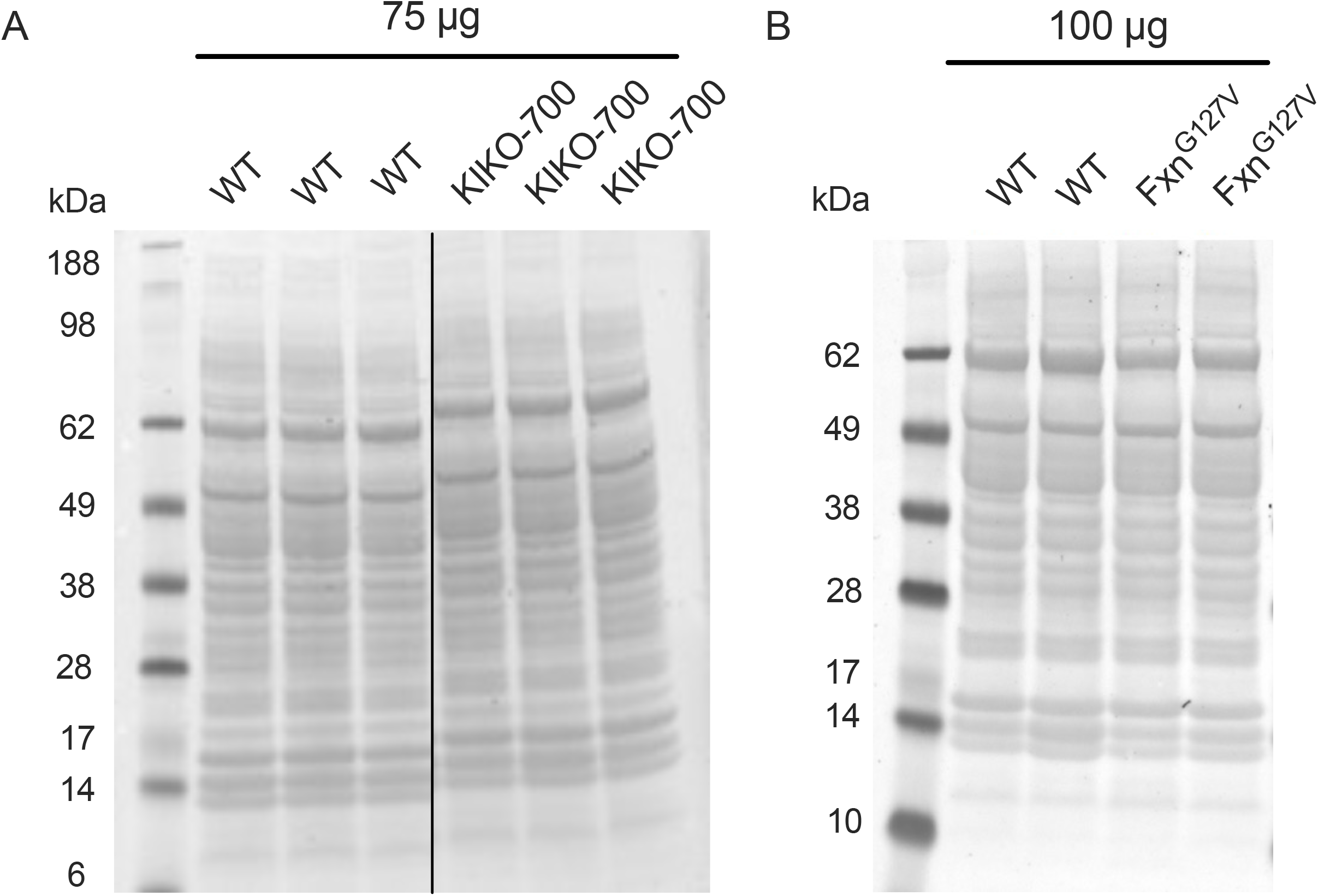
FXN normalization blots. (A) Ponceau S staining of the membrane for KIKO-700 samples in Fig 2A (n = 3) (B) Ponceau S staining of the membrane for Fxn^G127V^ samples, where 100 μg were loaded in Fig. 3A (n = 2).

**Figure S3.**
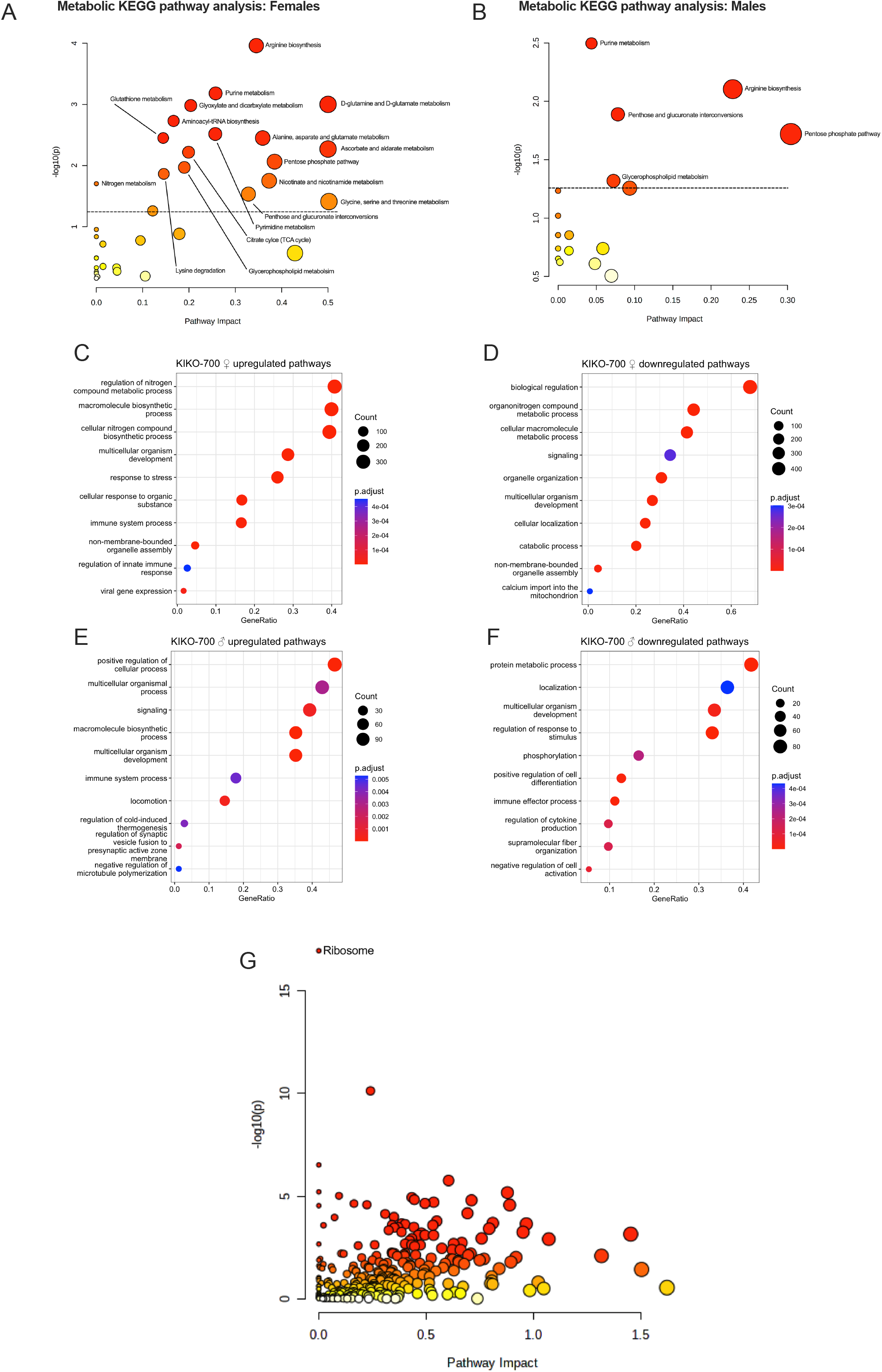
Enrichment of transcriptional and metabolic pathways associated with metabolic stress in KIKO-700 hearts. (A, B) Unbiased metabolic KEGG pathway analysis for (A) females and (B) males. (C, D) Top 10 results of unbiased GO analyses of significantly upregulated (C) or downregulated (D) genes in female KIKO-700 hearts. (E, F) Top 10 results of unbiased GO analyses of significantly upregulated (E) or downregulated (F) genes in male KIKO-700 hearts. (G) Integrated metabolomics and transcriptomics data from KIKO-700 females including ribosomal genes. n = 3/sex/genotype 18-months old.

**Figure S4.**
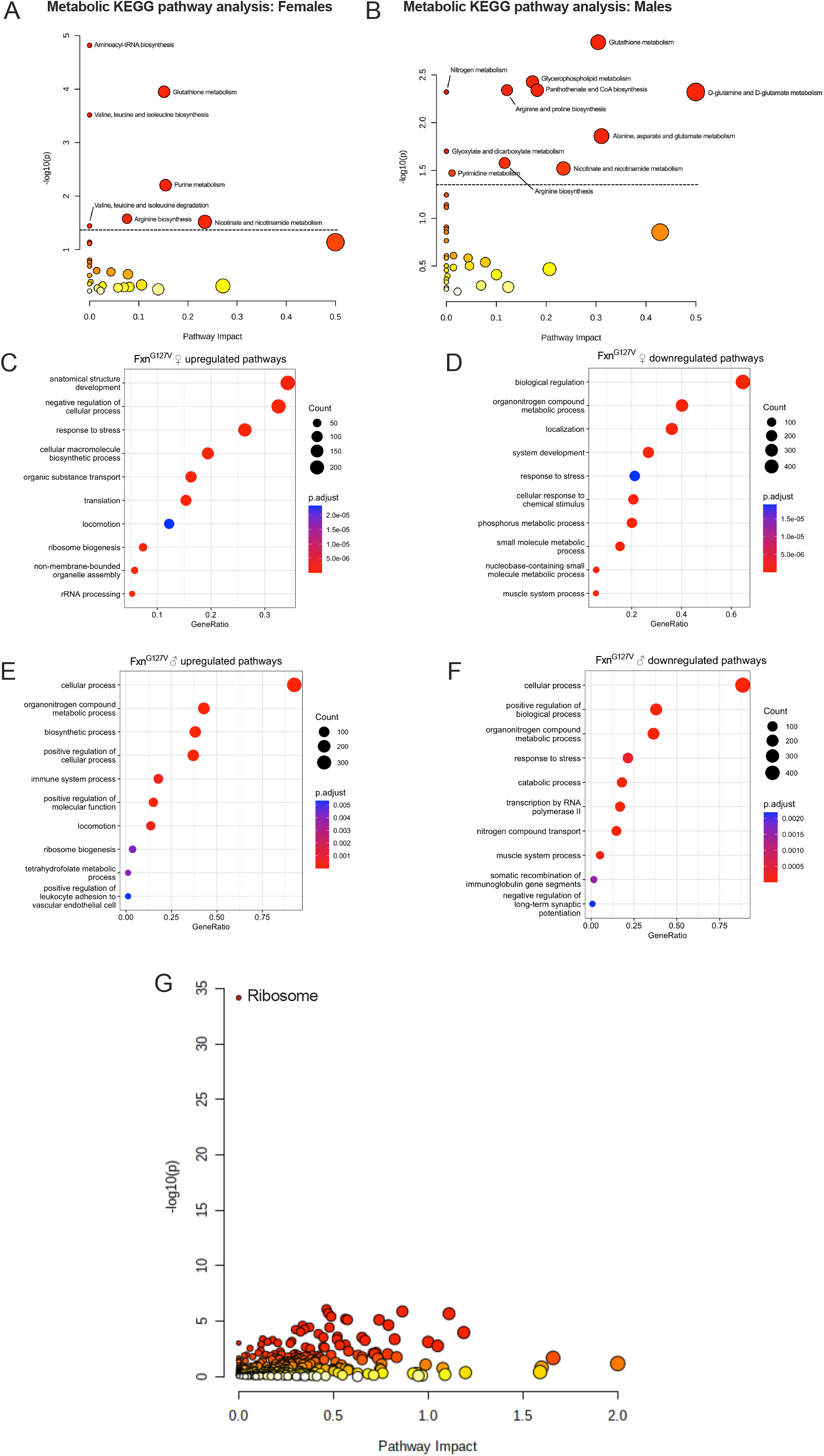
Enrichment of transcriptional and metabolic pathways associated with cardiac stress and ISR^mt^ in Fxn^G127V^ hearts. (A, B) Unbiased metabolic KEGG pathway analysis for (A) females and (B) males. (C, D) Top 10 results of unbiased GO analyses of significantly upregulated (C) or downregulated (D) genes in female Fxn^G127V^ hearts. (E, F) Top 10 results of unbiased GO analyses of significantly upregulated (E) or downregulated (F) genes in male Fxn^G127V^ hearts. (G) Integrated metabolomics and transcriptomics data from Fxn^G127V^ females including ribosomal genes. n = 3/sex/genotype 18-months old.

**Figure S5.**
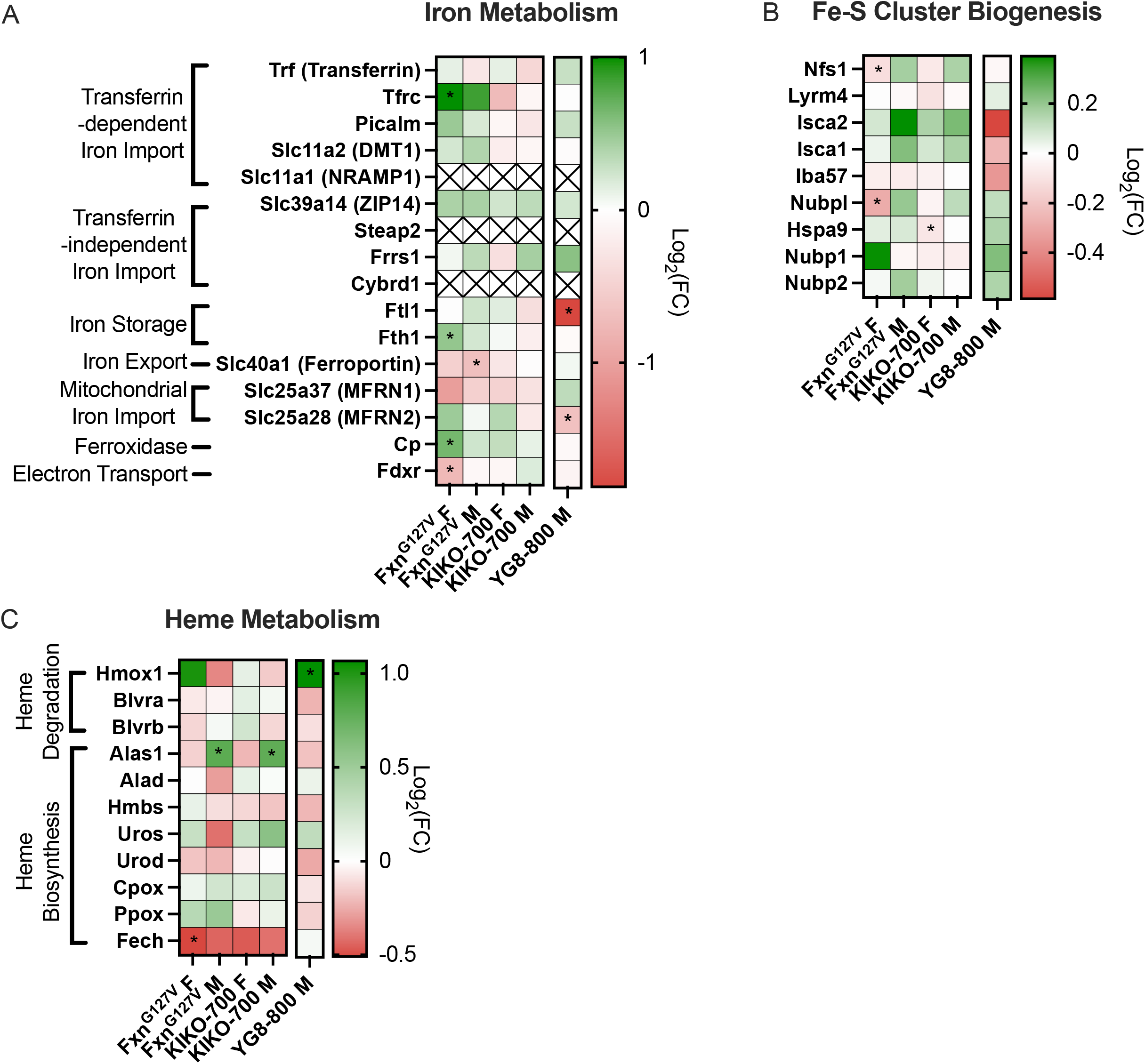
Lack of overall expression changes in iron metabolism genes in all three FXN-deficiency models. (A-C) Heatmap showing the expression of genes related to (A) iron metabolism, (B) Fe-S cluster biogenesis, and (C) heme metabolism. n = 3/sex/genotype 18-months old FXN^G127V^ and KIKO-800 and n = 4 18-months old YG8-800. * = p < 0.05.

**Figure S6.**
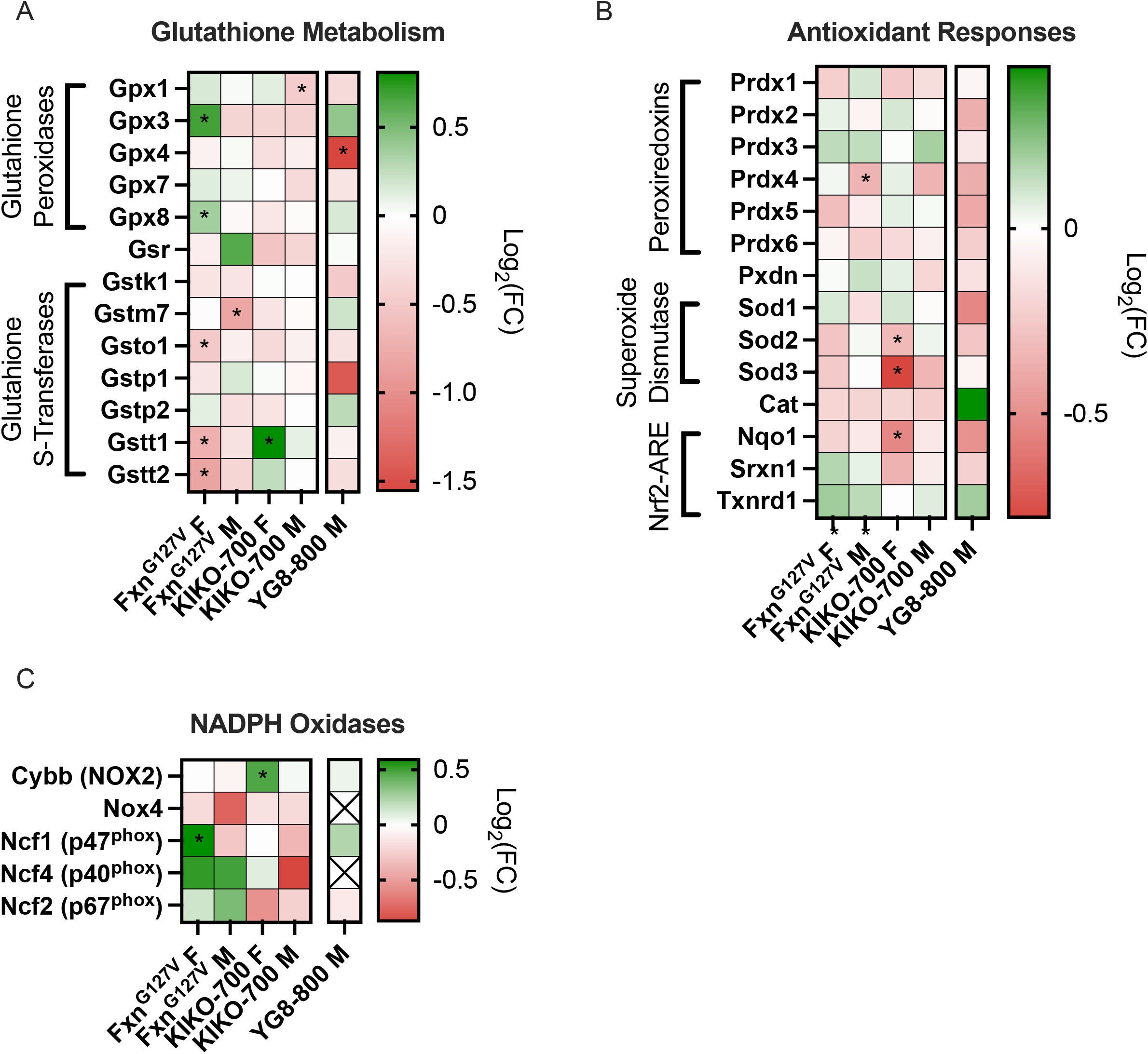
Absence of oxidative stress markers in all three FXN-deficiency models. (A-C) Heatmap showing the expression of genes related to (A) glutathione metabolism, (B) antioxidant responses, and (C) NADPH oxidases. n = 3/sex/genotype 18-months old FXN^G127V^ and KIKO-800 and n = 4 18-months old YG8-800. * = p < 0.05.

## References

Anjomani Virmouni, S., Ezzatizadeh, V., Sandi, C., Sandi, M., Al-Mahdawi, S., Chutake, Y. and Pook, M. A. (2015). A novel GAA-repeat-expansion-based mouse model of Friedreich’s ataxia. Dis Model Mech 8, 225–35.

Aramburu, J., Ortells, M. C., Tejedor, S., Buxade, M. and Lopez-Rodriguez, C. (2014). Transcriptional regulation of the stress response by mTOR. Sci Signal 7, re2.

Bidichandani, S. I., Ashizawa, T. and Patel, P. I. (1997). Atypical Friedreich ataxia caused by compound heterozygosity for a novel missense mutation and the GAA triplet-repeat expansion. Am J Hum Genet 60, 1251–6.

Bidichandani, S. I. and Delatycki, M. B. (1993). Friedreich Ataxia. In GeneReviews((R)), (eds M. P. Adam D. B. Everman G. M. Mirzaa R. A. Pagon S. E. Wallace L. J. H. Bean K. W. Gripp and A. Amemiya). Seattle (WA).

Boenzi, S. and Diodato, D. (2018). Biomarkers for mitochondrial energy metabolism diseases. Essays Biochem 62, 443–454.

Campuzano, V., Montermini, L., Molto, M. D., Pianese, L., Cossee, M., Cavalcanti, F., Monros, E., Rodius, F., Duclos, F., Monticelli, A. et al. (1996). Friedreich’s ataxia: autosomal recessive disease caused by an intronic GAA triplet repeat expansion. Science 271, 1423–7.

Chen, F., Haigh, S., Barman, S. and Fulton, D. J. (2012). From form to function: the role of Nox4 in the cardiovascular system. Front Physiol 3, 412.

Chiang, S., Braidy, N., Maleki, S., Lal, S., Richardson, D. R. and Huang, M. L. (2021). Mechanisms of impaired mitochondrial homeostasis and NAD(+) metabolism in a model of mitochondrial heart disease exhibiting redox active iron accumulation. Redox Biol 46, 102038.

Dogan, S. A., Pujol, C., Maiti, P., Kukat, A., Wang, S., Hermans, S., Senft, K., Wibom, R., Rugarli, E. I. and Trifunovic, A. (2014). Tissue-specific loss of DARS2 activates stress responses independently of respiratory chain deficiency in the heart. Cell Metab 19, 458–69.

Eckl, E. M., Ziegemann, O., Krumwiede, L., Fessler, E. and Jae, L. T. (2021). Sensing, signaling and surviving mitochondrial stress. Cell Mol Life Sci 78, 5925–5951.

Fil, D., Chacko, B. K., Conley, R., Ouyang, X., Zhang, J., Darley-Usmar, V. M., Zuberi, A. R., Lutz, C. M., Napierala, M. and Napierala, J. S. (2020). Mitochondrial damage and senescence phenotype of cells derived from a novel frataxin G127V point mutation mouse model of Friedreich’s ataxia. Dis Model Mech 13.

Forsstrom, S., Jackson, C. B., Carroll, C. J., Kuronen, M., Pirinen, E., Pradhan, S., Marmyleva, A., Auranen, M., Kleine, I. M., Khan, N. A. et al. (2019). Fibroblast Growth Factor 21 Drives Dynamics of Local and Systemic Stress Responses in Mitochondrial Myopathy with mtDNA Deletions. Cell Metab 30, 1040–1054 e7.

Galea, C. A., Huq, A., Lockhart, P. J., Tai, G., Corben, L. A., Yiu, E. M., Gurrin, L. C., Lynch, D. R., Gelbard, S., Durr, A. et al. (2016). Compound heterozygous FXN mutations and clinical outcome in friedreich ataxia. Ann Neurol 79, 485–95.

Gerard, C., Archambault, A. F., Bouchard, C. and Tremblay, J. P. (2023). A promising mouse model for Friedreich Ataxia progressing like human patients. Behav Brain Res 436, 114107.

Hanson, E., Sheldon, M., Pacheco, B., Alkubeysi, M. and Raizada, V. (2019). Heart disease in Friedreich’s ataxia. World J Cardiol 11, 1–12.

Huang, M. L., Sivagurunathan, S., Ting, S., Jansson, P. J., Austin, C. J., Kelly, M., Semsarian, C., Zhang, D. and Richardson, D. R. (2013). Molecular and functional alterations in a mouse cardiac model of Friedreich ataxia: activation of the integrated stress response, eIF2alpha phosphorylation, and the induction of downstream targets. Am J Pathol 183, 745–57.

Isnard, R., Kalotka, H., Durr, A., Cossee, M., Schmitt, M., Pousset, F., Thomas, D., Brice, A., Koenig, M. and Komajda, M. (1997). Correlation between left ventricular hypertrophy and GAA trinucleotide repeat length in Friedreich’s ataxia. Circulation 95, 2247–9.

Kaspar, S., Oertlin, C., Szczepanowska, K., Kukat, A., Senft, K., Lucas, C., Brodesser, S., Hatzoglou, M., Larsson, O., Topisirovic, I. et al. (2021). Adaptation to mitochondrial stress requires CHOP-directed tuning of ISR. Sci Adv 7.

Khan, N. A., Nikkanen, J., Yatsuga, S., Jackson, C., Wang, L., Pradhan, S., Kivela, R., Pessia, A., Velagapudi, V. and Suomalainen, A. (2017). mTORC1 Regulates Mitochondrial Integrated Stress Response and Mitochondrial Myopathy Progression. Cell Metab 26, 419–428 e5.

Koeppen, A. H. (2011). Friedreich’s ataxia: pathology, pathogenesis, and molecular genetics. J Neurol Sci 303, 1–12.

Kolberg, L., Raudvere, U., Kuzmin, I., Vilo, J. and Peterson, H. (2020). gprofiler2 -- an R package for gene list functional enrichment analysis and namespace conversion toolset g:Profiler. F1000Res 9.

Kuhl, I., Miranda, M., Atanassov, I., Kuznetsova, I., Hinze, Y., Mourier, A., Filipovska, A. and Larsson, N. G. (2017). Transcriptomic and proteomic landscape of mitochondrial dysfunction reveals secondary coenzyme Q deficiency in mammals. Elife 6.

Lamarche, J. B., Cote, M. and Lemieux, B. (1980). The cardiomyopathy of Friedreich’s ataxia morphological observations in 3 cases. Can J Neurol Sci 7, 389–96.

Li, J., Rozwadowska, N., Clark, A., Fil, D., Napierala, J. S. and Napierala, M. (2019). Excision of the expanded GAA repeats corrects cardiomyopathy phenotypes of iPSC-derived Friedreich’s ataxia cardiomyocytes. Stem Cell Res 40, 101529.

Li, Y., Polak, U., Clark, A. D., Bhalla, A. D., Chen, Y. Y., Li, J., Farmer, J., Seyer, L., Lynch, D., Butler, J. S. et al. (2016). Establishment and Maintenance of Primary Fibroblast Repositories for Rare Diseases-Friedreich’s Ataxia Example. Biopreserv Biobank 14, 324–9.

Liao, Y., Smyth, G. K. and Shi, W. (2019). The R package Rsubread is easier, faster, cheaper and better for alignment and quantification of RNA sequencing reads. Nucleic Acids Res 47, e47.

Lopaschuk, G. D. and Jaswal, J. S. (2010). Energy metabolic phenotype of the cardiomyocyte during development, differentiation, and postnatal maturation. J Cardiovasc Pharmacol 56, 130–40.

Lopaschuk, G. D., Ussher, J. R., Folmes, C. D., Jaswal, J. S. and Stanley, W. C. (2010). Myocardial fatty acid metabolism in health and disease. Physiol Rev 90, 207–58.

Martelli, A. and Puccio, H. (2014). Dysregulation of cellular iron metabolism in Friedreich ataxia: from primary iron-sulfur cluster deficit to mitochondrial iron accumulation. Front Pharmacol 5, 130.

Martelli, A., Schmucker, S., Reutenauer, L., Mathieu, J. R. R., Peyssonnaux, C., Karim, Z., Puy, H., Galy, B., Hentze, M. W. and Puccio, H. (2015). Iron regulatory protein 1 sustains mitochondrial iron loading and function in frataxin deficiency. Cell Metab 21, 311–323.

Mehrmohamadi, M., Liu, X., Shestov, A. A. and Locasale, J. W. (2014). Characterization of the usage of the serine metabolic network in human cancer. Cell Rep 9, 1507–19.

Miranda, C. J., Santos, M. M., Ohshima, K., Smith, J., Li, L., Bunting, M., Cossee, M., Koenig, M., Sequeiros, J., Kaplan, J. et al. (2002). Frataxin knockin mouse. FEBS Lett 512, 291–7.

Morgan, M., Anders, S., Lawrence, M., Aboyoun, P., Pages, H. and Gentleman, R. (2009). ShortRead: a bioconductor package for input, quality assessment and exploration of high-throughput sequence data. Bioinformatics 25, 2607–8.

Nabeebaccus, A. A., Zoccarato, A., Hafstad, A. D., Santos, C. X., Aasum, E., Brewer, A. C., Zhang, M., Beretta, M., Yin, X., West, J. A. et al. (2017). Nox4 reprograms cardiac substrate metabolism via protein O-GlcNAcylation to enhance stress adaptation. JCI Insight 2.

Narayana Rao, K. B., Pandey, P., Sarkar, R., Ghosh, A., Mansuri, S., Ali, M., Majumder, P., Ranjith Kumar, K., Ray, A., Raychaudhuri, S. et al. (2022). Stress Responses Elicited by Misfolded Proteins Targeted to Mitochondria. J Mol Biol 434, 167618.

Nikkanen, J., Forsstrom, S., Euro, L., Paetau, I., Kohnz, R. A., Wang, L., Chilov, D., Viinamaki, J., Roivainen, A., Marjamaki, P. et al. (2016). Mitochondrial DNA Replication Defects Disturb Cellular dNTP Pools and Remodel One-Carbon Metabolism. Cell Metab 23, 635–48.

O’Connell, T. M., Logsdon, D. L. and Payne, R. M. (2022). Metabolomics analysis reveals dysregulation in one carbon metabolism in Friedreich Ataxia. Mol Genet Metab 136, 306–314.

Palau, F. (2001). Friedreich’s ataxia and frataxin: molecular genetics, evolution and pathogenesis (Review). Int J Mol Med 7, 581–9.

Payne, R. M. (2022). Cardiovascular Research in Friedreich Ataxia: Unmet Needs and Opportunities. JACC Basic Transl Sci 7, 1267–1283.

Payne, R. M. and Wagner, G. R. (2012). Cardiomyopathy in Friedreich ataxia: clinical findings and research. J Child Neurol 27, 1179–86.

Perdomini, M., Belbellaa, B., Monassier, L., Reutenauer, L., Messaddeq, N., Cartier, N., Crystal, R. G., Aubourg, P. and Puccio, H. (2014). Prevention and reversal of severe mitochondrial cardiomyopathy by gene therapy in a mouse model of Friedreich’s ataxia. Nat Med 20, 542–7.

Puccio, H. and Koenig, M. (2000). Recent advances in the molecular pathogenesis of Friedreich ataxia. Hum Mol Genet 9, 887–92.

Puccio, H., Simon, D., Cossee, M., Criqui-Filipe, P., Tiziano, F., Melki, J., Hindelang, C., Matyas, R., Rustin, P. and Koenig, M. (2001). Mouse models for Friedreich ataxia exhibit cardiomyopathy, sensory nerve defect and Fe-S enzyme deficiency followed by intramitochondrial iron deposits. Nat Genet 27, 181–6.

Ritchie, M. E., Phipson, B., Wu, D., Hu, Y., Law, C. W., Shi, W. and Smyth, G. K. (2015). limma powers differential expression analyses for RNA-sequencing and microarray studies. Nucleic Acids Res 43, e47.

Rotig, A., de Lonlay, P., Chretien, D., Foury, F., Koenig, M., Sidi, D., Munnich, A. and Rustin, P. (1997). Aconitase and mitochondrial iron-sulphur protein deficiency in Friedreich ataxia. Nat Genet 17, 215–7.

Sasagawa, S., Nishimura, Y., Okabe, S., Murakami, S., Ashikawa, Y., Yuge, M., Kawaguchi, K., Kawase, R., Okamoto, R., Ito, M. et al. (2016). Downregulation of GSTK1 Is a Common Mechanism Underlying Hypertrophic Cardiomyopathy. Front Pharmacol 7, 162.

Sasset, L., Manzo, O. L., Zhang, Y., Marino, A., Rubinelli, L., Riemma, M. A., Chalasani, M. L. S., Dasoveanu, D. C., Roviezzo, F., Jankauskas, S. S. et al. (2022). Nogo-A reduces ceramide de novo biosynthesis to protect from heart failure. Cardiovasc Res.

Sayles, N. M., Southwell, N., McAvoy, K., Kim, K., Pesini, A., Anderson, C. J., Quinzii, C., Cloonan, S., Kawamata, H. and Manfredi, G. (2022). Mutant CHCHD10 causes an extensive metabolic rewiring that precedes OXPHOS dysfunction in a murine model of mitochondrial cardiomyopathy. Cell Rep 38, 110475.

Smyrnias, I. (2021). The mitochondrial unfolded protein response and its diverse roles in cellular stress. Int J Biochem Cell Biol 133, 105934.

Tong, W. H., Ollivierre, H., Noguchi, A., Ghosh, M. C., Springer, D. A. and Rouault, T. A. (2022). Hyperactivation of mTOR and AKT in a cardiac hypertrophy animal model of Friedreich ataxia. Heliyon 8, e10371.

Vasquez-Trincado, C., Patel, M., Sivaramakrishnan, A., Bekeova, C., Anderson-Pullinger, L., Wang, N., Tang, H. Y. and Seifert, E. L. (2021). Adaptation of the heart to Frataxin depletion: Evidence that integrated stress response can predominate over mTORC1 activation. Hum Mol Genet.

Vigil-Garcia, M., Demkes, C. J., Eding, J. E. C., Versteeg, D., de Ruiter, H., Perini, I., Kooijman, L., Gladka, M. M., Asselbergs, F. W., Vink, A. et al. (2021). Gene expression profiling of hypertrophic cardiomyocytes identifies new players in pathological remodelling. Cardiovasc Res 117, 1532–1545.

Wu, C. W. and Storey, K. B. (2021). mTOR Signaling in Metabolic Stress Adaptation. Biomolecules 11.

Wu, T., Hu, E., Xu, S., Chen, M., Guo, P., Dai, Z., Feng, T., Zhou, L., Tang, W., Zhan, L. et al. (2021). clusterProfiler 4.0: A universal enrichment tool for interpreting omics data. Innovation (Camb) 2, 100141.

Xia, J., Psychogios, N., Young, N. and Wishart, D. S. (2009). MetaboAnalyst: a web server for metabolomic data analysis and interpretation. Nucleic Acids Res 37, W652–60.

Zhao, Q. D., Viswanadhapalli, S., Williams, P., Shi, Q., Tan, C., Yi, X., Bhandari, B. and Abboud, H. E. (2015). NADPH oxidase 4 induces cardiac fibrosis and hypertrophy through activating Akt/mTOR and NFkappaB signaling pathways. Circulation 131, 643–55.

